# Simulating structurally variable Nuclear Pore Complexes for Microscopy

**DOI:** 10.1101/2022.05.17.492295

**Authors:** Maria Theiss, Jean-Karim Hériché, Craig Russell, David Helekal, Alisdair Soppitt, Jonas Ries, Jan Ellenberg, Alvis Brazma, Virginie Uhlmann

## Abstract

**Motivation:** The Nuclear Pore Complex (NPC) is the only passageway for macromolecules between nucleus and cytoplasm, and one of localization microscopy’s most important reference standards: it is massive and stereotypically arranged. The average architecture of NPC proteins has been resolved with pseudo-atomic precision, however observed NPC heterogeneities evidence a high degree of divergence from this average. Single Molecule Localization Microscopy (SMLM) images NPCs at protein-level resolution, whereupon image analysis software studies NPC variability. However the true picture of NPC variability is unknown. In quantitative image analysis experiments, it is thus difficult to distinguish intrinsically high SMLM noise from true variability of the underlying structure.

**Results:** We introduce CIR4MICS (“ceramics”, Configurable, Irregular Rings FOR MICroscopy Simulations), a pipeline that creates artificial datasets of structurally variable synthetic NPCs based on architectural models of the true NPC. Users can select one or more N- or C-terminally tagged NPC proteins, and simulate a wide range of geometric variations. We also represent the NPC as a spring-model such that arbitrary deforming forces, of user-defined magnitudes, simulate irregularly shaped variations. We provide an open-source simulation pipeline, as well as reference datasets of simulated human NPCs. Accompanying ground truth annotations allow to test the capabilities of image analysis software and facilitate a side-by-side comparison with real data. We demonstrate this by synthetically replicating a geometric analysis of real NPC radii and reveal that a wide range of simulated variability parameters can lead to observed results. Our simulator is therefore valuable to benchmark and develop image analysis methods, as well as to inform experimentalists about the requirements of hypothesis-driven imaging studies.

**Availability:** Code: https://github.com/uhlmanngroup/cir4mics. Simulated data is available at BioStudies (Accession number S-BSST1058).

**Supplementary information:** Supplementary data are available at

## Introduction

Nuclear Pore Complexes (NPCs) enable all macromolecular exchange between cytoplasm and nucleus (Paci, Caria, and Lemke, 2021). The human NPC consists of ∼30 types of proteins, called Nucleoporins (Nups) (Reichelt, 1990; Schwartz, 2016), which arrange eight-fold symmetrically around a transport axis, and form three stacked rings along the Nuclear envelope (NE) plane: the nucleoplasmic ring (NR) and cytoplasmic ring (CR) are called outer rings (OR) as they sandwich an inner ring (IR) (Lin and Hoelz, 2019) (Figure 1, left; Figure 2). The NPC’s abundance, size, and geometry has rendered it one of super-resolution microscopy’s most important reference standard (Thevathasan et al., 2019; Mund and Ries, 2020; Sage et al., 2019).

**Fig. 1.**
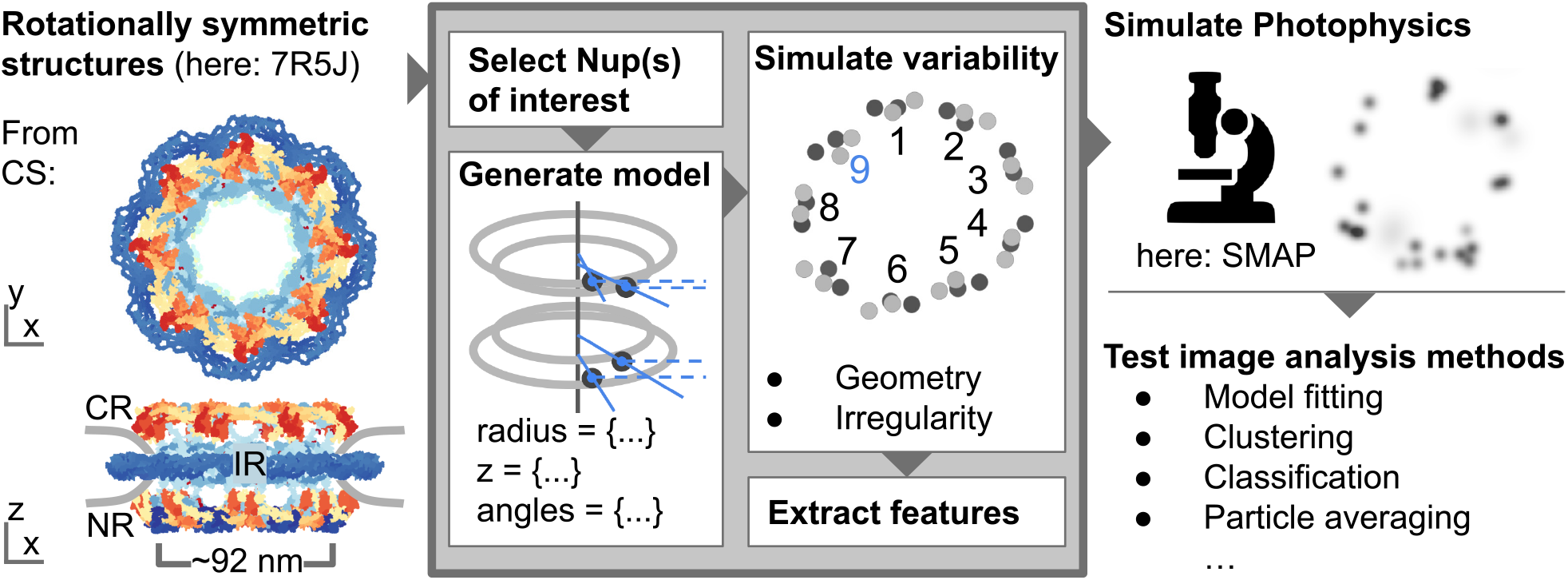
CIR4MICS workflow — our work is boxed. Left: rotationally symetric PDB models function as input, 5A9Q (Appen et al., 2015); 7PEQ, 7PER (Schuller et al., 2021); 5IJN, 5IJO (Kosinski et al., 2016); 7R5J, 7R5K (Mosalaganti et al., 2022). Model 7R5J (Mosalaganti et al., 2022) is shown as an example. left, top: NPC in top-view from the cytoplasmic side (CS). NE is not shown. Left bottom: NPC in side-view with cytoplasmic ring (CR), Inner ring (IR), and Nucleoplasmic ring (NR). Nuclear envelope is shown in cut side-view overlaid by luminal Nup210 (dark blue). 7R5J represents the NPC in intact cells with membrane-pore diameter of 92 nm. Boxed, centre: One or more Nups can be selected for labelling. Each individual Nup is encoded with a radius, angle, and z-position. Geometric or irregularly shaped variability can be added and ground-truth features are extracted. Right: Coordinates are passed to microscopy simulation software (e.g. Ries (2020)). Raw coordinates, photophysics representation, and ground truth features can be used to benchmark Image analysis software.

**Fig. 2.**
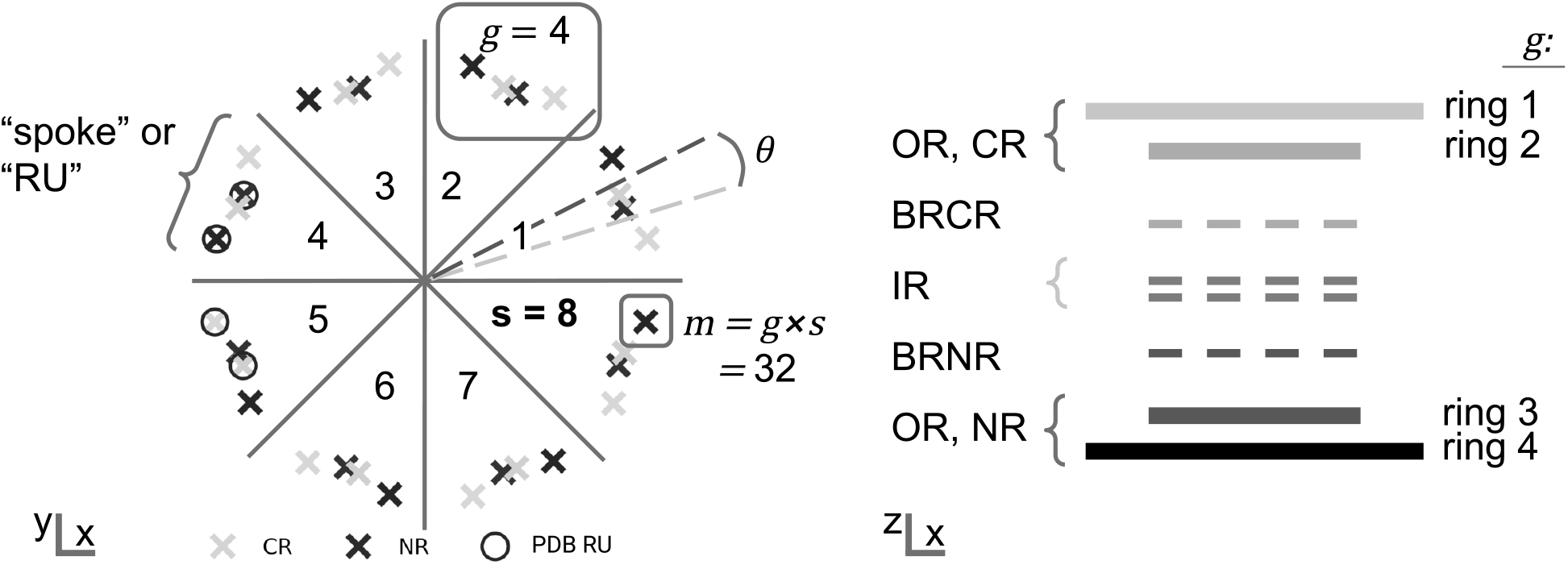
Layout of the spring model: Position of an example Nup (N terminus of Nup107 in PDB 5A9Q, Appen et al. (2015)) in top view (left) and side view (right, solid lines only). Nup107 occurs in *g* = 4 rings and has symmetry *s* = 8, corresponding to a total copy number of *m* = *gs* = 32. A RU according to the PDB assembly is indicated (PDB RU). The PDB RU can be modified to our RU by offsetting the CR or NR by *±*2*π/s* such that their twist angle *θ* is the most acute (Suppl. section 7.4). Right: Difference between OR, NR, CR, IR, BRCR, BRNR, and “ring”.

This structure has been solved with pseudo-atomic resolution (Appen et al., 2015; Kosinski et al., 2016; Schuller et al., 2021). Recent breakthroughs in *in silico* protein-structure prediction enabled near-complete models of the NPC scaffold (Jumper et al., 2021; Mosalaganti et al., 2022). However, these models achieve their high resolution by integrating information from multiple sources, such as Electron Microscopy (EM), Nup atomic structures, and protein interaction studies, averaging out hundreds of NPCs and their variability (Rantos, Karius, and Kosinski, 2021).

Examples of NPC variability are deviations from the 8-fold rotational symmetry (Hinshaw and Milligan, 2003; Löschberger et al., 2014; Stanley, Fassati, and Hoogenboom, 2018), elliptical deformations (Beck et al., 2007; Maimon et al., 2012; Huijben et al., 2021), varying azimuthal “twist” angle between CR and NR (Heydarian et al., 2021), and varying pore diameters (Stanley, Fassati, and Hoogenboom, 2018; Sabinina et al., 2021; Zimmerli et al., 2021; Mosalaganti et al., 2022). Furthermore, the distance between CR and NR is inconsistently reported (Sabinina et al., 2021; Heydarian et al., 2021; Curd et al., 2020). This natural NPC variability remains to be fully characterised. To this end, coarse-grained models and coarse grained elastic network models predicted possible deformations of the yeast NPC (Wolf and Mofrad, 2008; Lezon, Sali, and Bahar, 2009). Molecular dynamics have been used to simulate lateral tension on the NE, which dilated a model NPC (Mosalaganti et al., 2022). Theoretical models aim to predict NPC heterogeneity and are often motivated by imaging studies (Zimmerli et al., 2021; Mosalaganti et al., 2022), but conversely, can form hypotheses to be tested *ex silico*.

For example Atomic Force Microscopy and EM have been used to study NPC shape variability (e.g. Stanley, Fassati, and Hoogenboom, 2018; Hinshaw and Milligan, 2003). Both microscopy techniques however cannot elucidate the interplay of different Nups of interest (NoIs) in structurally variable NPCs, as neither commonly tag specific NoI, and both lack the resolution to discern NoI at the single pore level. Fluorescence microscopy on the other hand can specifically tag NoIs (Li et al., 2022), however the diffraction limit of visible light is typically double the size of a human NPC (Abbe, 1873; Mosalaganti et al., 2022).

Single-Molecule Localisation Microscopy (SMLM) is a fluorescence microscopy technique that circumvents the diffraction limit and can therefore be used to analyse NPC heterogeneity at single-pore level and for specific Nups (Wu, Hoess, et al., 2022; Huijben et al., 2021; Sabinina et al., 2021; Wu, Tschanz, et al., 2020). More precisely, fluorophores in SMLM blink stochastically such that only one fluorophore within a neighbourhood is active at a time; it is thus resolvable from its neighbours (Rust, Bates, and Zhuang, 2006). SMLM has been shown to image 4 differently labelled Nups in 3D (Testa et al., 2010; Li et al., 2022; Huang et al., 2008; Middendorff et al., 2008) and to resolve Nups that are 12 nm apart (Diekmann et al., 2020; Thevathasan et al., 2019). This results in a high-contrast, sparse, coordinate representation of each NPC, which can be fitted, classified, or averaged by downstream software (Wu, Hoess, et al., 2022; Sabinina et al., 2021; Huijben et al., 2021).

SMLM however is prone to noise, aberrations, and artefacts. (Jacquemet et al., 2020; Thevathasan et al., 2019). Consequently, it is near impossible to automatically or manually annotate structural variability of individual pores with high confidence. The ability of software to analyse NPC heterogeneity cannot be assessed without ground truth, whereas ground truth cannot be assigned without validated computational analysis. Simulated NPCs with known variability can solve this catch-22.

Wu, Hoess, et al. (2022) simulate a model of a Nup96 that can be varied in its radius, distance between the ORs, ellipticity, and twist angle between the ORs. Multiple approaches have been developed to simulate noise throughout the SMLM imaging pipeline, for instance sparse labelling and linkage error, and detector readout noise (Sinkó et al., 2014; Venkataramani et al., 2016; Novák et al., 2017; Griffié et al., 2020; Sage et al., 2019; Ries, 2020). Wu, Hoess, et al. (2022) and Ries (2020) simulate non-geometric shape variations as part of imaging noise by randomly and independently offsetting localisations from their initial positions.

These approaches however do not explicitly simulate irregular NPC shapes, i.e. shapes that are coherent, yet non-geometric, as visible in imaging studies (Stanley, Fassati, and Hoogenboom, 2018; Beton et al., 2021; Löschberger et al., 2014). Existing simulations furthermore only integrate a very limited pre-selection of Nups (typically one), integrated Nups are only loosely derived from real NPCs (Huijben et al., 2021; Venkataramani et al., 2016; Wu, Hoess, et al., 2022), and often 2D (Huijben et al., 2021; Venkataramani et al., 2016). However, explicit simulations of NPC shape irregularity are paramount to study the robustness of fitting, classification, and averaging software to hypothesised shape irregularity. Simulations of this kind enable the development and benchmarking of software applications that can distinguish between imaging noise, shape irregularity, and geometry, thereby shedding light on NPC biology.

CIR4MICS includes a computationally light-weight spring based model of the NPC to simulate a wide range of geometric variations and irregularly shaped variations for SMLM. We provide a wide range of Nup models for selection which are N- and C-terminally labelled, and from human NPCs in either NE isolates or intact cells (Appen et al., 2015; Schuller et al., 2021; Kosinski et al., 2016; Mosalaganti et al., 2022). Nups can be chosen for both single-channel and multi-channel simulations, as shape irregularity is coherent across all selected types of Nups. Additionally, we provide datasets of geometrically variable and irregularly shaped NPCs, which includes simulated SMLM data using existing software (Ries, 2020) together with our underlying ground truth coordinates and features. CIR4MICS offers a framework for benchmarking image analysis methods that are general purpose, or NPC specific, the latter of which is helpful to elucidate analysis software’s ability to capture NPC biological variability.

## Methods

### Spring models simulate irregular but coherent deformations

We simulate the NPC as a partially connected spring model to which we apply deforming forces. This results in shape variations that can be irregular due to the stochasticity of deforming forces, yet coherent as springs ensure forces are distributed throughout the model (Figure S1A). These simulations are performed on individual types of Nups, however we use “NPC” as an umbrella term for any Nup of interest (NoI). Springs represent the elastic properties of proteins and membranes, whereby forces represent any cause why the NPC is not observed in its perfectly geometrical shape. Upon application of a force, springs will be displaced from their resting lengths and oscillate to a new equilibrium. If oscillatory energy dissipates, and if the applied force is static, the new equilibrium will also be static. SMLM has a low temporal resolution and commonly images fixed cells. SMLM data that resolves NPC shape variability take minutes to acquire (Thevathasan et al., 2019; Diekmann et al., 2020). These data do not temporally resolve the microsecond-to second-long nucleocytoplasmic transport events (Kubitscheck and Siebrasse, 2017). To simulate NPCs, we therefore damp spring oscillation, and keep forces static. Consequently, we only use the static new equilibrium for downstream SMLM simulations of NPCs.

### NPC structural models function as input

Pseudo-atomic structural models of the NPC exceed the resolution of SMLM and contain multiple types of Nups, facilitating multi-colour simulations. SMLM usually captures NPCs in whole, but chemically fixed cells. However, models 5A9Q (Appen et al., 2015), 5IJN, 5IJO (Kosinski et al., 2016), and 7R5K (Mosalaganti et al., 2022) represent NPCs in isolated NEs, whereas 7PEQ, 7PER (Schuller et al., 2021), and 7R5J (Mosalaganti et al., 2022) represent NPCs in their native environment. It is therefore uncertain which models best represent NPCs under SMLM conditions. Furthermore, live-cell SMLM, in which NPCs are in their native environment, might become more common in the future (Lincoln et al., 2021). We therefore made all mentioned human NPC models available in our framework (Table S2 and Table S3). Our pipeline also allows for the addition of further models (Figure 1).

### Implementing NPC structural models

#### Nomenclature

Nups within the NPCs in top-view are arranged in *s* symmetrical “spokes” around a central channel, where *s* = 8 is the default rotational symmetry of the NPC. “Rotational unit” (RU) refers to all structurally resolved Nups in one of *s* spokes. The copy number *g* of one NoI in one RU corresponds to its overall copy number *m* via *m* = *gs*. In side-view, it is accordingly arranged in *g* stacked “rings” that are parallel to the NE plane. The term “rings” is not to be confused with the terms “CR”, “NR”, or “IR”, as each of the latter three might contain several “rings” of a specific Nup (Figure 2). We furthermore denote an individual Nup as *p*, and a type of Nup as *P*. Important notation is summarised in Table S1.

#### Multi-colour simulations

Multi-channel SMLM has imaged four distinct types of Nups (Li et al., 2022). We therefore enable simulations with up to six labels. Distinct types of Nups can either be treated identically, or the first listed Nup type can be set as reference. Setting a reference Nup allows to compare its geometry between single- and multi-channel simulations.

#### Extracting Nup coordinates from PDB

All used PDB models of the NPC are rotationally symmetric. Hence, one RU represents the entire model given the axis of symmetry 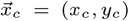 (Suppl. Section 2.1), and the order of symmetry *s*. Furthermore, protein termini are common fluorophore attachment sites, thus for each Nup within one rotational unit, we extract coordinates of the N-terminus and C-terminus. The user can specify up to six distinct labels, where each label consists of a type of Nup *P*_*i*_, *i* ∈ 1 … 6 and its terminus to be tagged. As most Nups occur in more than one copy per rotational unit, the total number *n* of fluorophore attachment sites with index *j* can be higher than the number of distinct labels. Each fluorophore attachment site is encoded via an axial position *z*_*j*_, radius *r*_*j*_ from 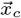, and an angle relative to other fluorophore attachment sites, *α*_*j*_ (Suppl. Section 2.2). The arithmetic mean *ā* of *α*_*j*_ is ffset to 0 for single-channel simulations. For multi-channel simulations, *ā* = 0 if no Nup *P*_*i*_ is selected as reference. Otherwise, *P*_1_ = *P*_ref_ with corresponding *ā*_ref_ and offset *ā* such that *ā*_ref_ = 0.

Values are converted from Å to nm, Cartesian coordinates are generated by reflecting coordinates of the RU *s* times around a new central axis 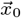 with 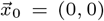 with an offset of 2*π/s* rad. Coordinates of each NoI are henceforth called “nodes”. A script that reads out and formats coordinates from models 7R5K and 7R5J (Mosalaganti et al., 2022) is available and can be adapted to other structural models.

### Layout of the spring model

The layout of the spring model is an abstraction of the NPC. We simulate each ring *g* as a separate spring model to ensure extensibility and comparability regardless of the selection and number of tagged Nup types. For each ring, nodes are anchored to the centre via radial springs (Figure 3A). These springs approximate the NE, whereby interior pressure replaces exterior contraction-resisting tension (Zimmerli et al., 2021). Nodes are connected to their 2*h, h* ∈ 0 … ⌊*s/*2⌋, neighbours via circumferential springs, which represent interactions within the NPC. *h* is 2 by default, as higher values increase computational cost, and lower values lead to overly jagged deformations (Figure 3A). Shorter circumferential springs are stiffer than longer circumferential springs, indicated by their spring constants *k*. Radial springs are less stiff than the shortest circumferential spring, as we assume protein-protein interactions are stronger than protein-membrane interactions. (Suppl. section 3).

**Fig. 3.**
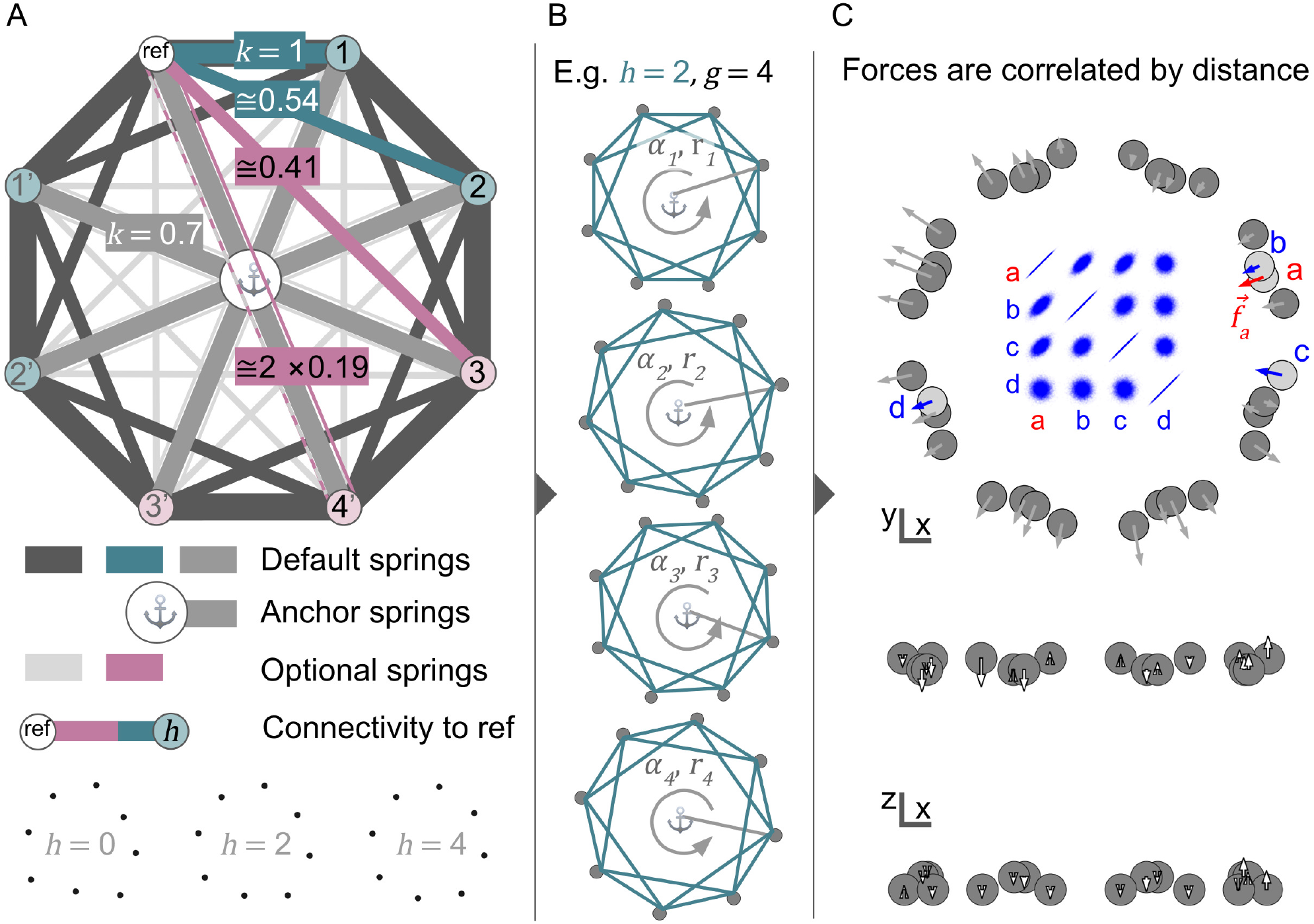
Layout of the spring model. **B:** Spring connectivity of one of *g* rings. By default, circumferential springs connect each node to their nearest *h* = 2 neighbours in clockwise (cw) and anticlockwise (acw) direction. cw default springs are shown in dark gray, and in teal for a reference node “ref”. Optional cw circumferential springs for ref, with *h* = 3 and *h* = 4, are shown in magenta. Spring constants *k* are shown on top of example springs and are decreasing with circumferential spring length. *h* = 4 connects opposing nodes with two springs whose spring-constant is halved to be equivalent to a single spring. All nodes are connected to a central anchor via springs with *k* = 0.7. Bottom: Effect of deformation *D*_mag_ 15 on nodes with *h* = 0, 2, 4. **C:** Each ring *g* is simulated separately and is defined by its radius *r*, rotation *α*, and *z*-position (*z* not shown). **D:** Continuity between independent rings is ensured with deforming forces (in x-y) or offsets (in z) that are covariant by distance. Top: 10000 forces (scalar values) sampled on exemplary nodes a, b, c, and d. Bottom: Axial offset (white arrows) are applied without springs.

### Stochastic forces deform the spring model laterally

Random, static forces are applied to nodes to achieve irregular shapes. Forces are sampled from a multivariate normal distribution with covariance dependent on the distance between nodes. Consequently, forces on nearby nodes are more similar than forces on distant nodes (Figure 3C). This ensures continuity between separate NPC rings and considers that forces on real NPCs are likely not pointforces. More precisely, covariance between forces on nodes 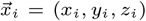 and 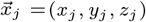, with *i, j* ∈ 1 … *m* are given by the RBF kernel,

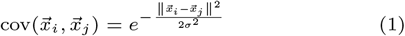

where, ∥·∥^2^ is the squared Euclidean norm, and *σ*is a free parameter. We couple *σ*to the size of the model by setting it to a fraction of the mean radius 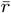. In comparison to a linear covariance between forces and distance, the RBF kernel ensures more similarity between forces on nearby nodes, preserving local structures. This is particularly important for nearby nodes on different rings. Simultaneously, forces on distant nodes are less similar, allowing for more irregular deformations. Empirically, 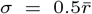 produces NPC deformations that are coherent at a short scale yet overall irregular (Figure S1B).

In addition, radii *r*_*i*_ of all nodes are transiently set to the arithmetic mean radius 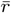 of all rings. Radial distances are therefore equalised, which preserves concentricity of rings during deformation.

Lastly, the magnitude of deformation *D*_mag_ is user-definable, so that the effects of different levels of deformation can be studied (Figure 4A, Figure S1B). *D*_mag_ [nm] covers ∼99.7 % of the sampling range (= 3 standard deviations = 3*σ*). It is then converted to variance *σ*^*′*2^ via *σ*^*′*2^ = (*D*_mag_*/*3)^2^. Next, cov is updated as cov^*′*^ = *σ*^*′*2^cov and scalar forces *f*_*i*_ with *i* ∈ 1 … *m* are sampled from

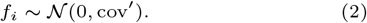

**Fig. 4.**
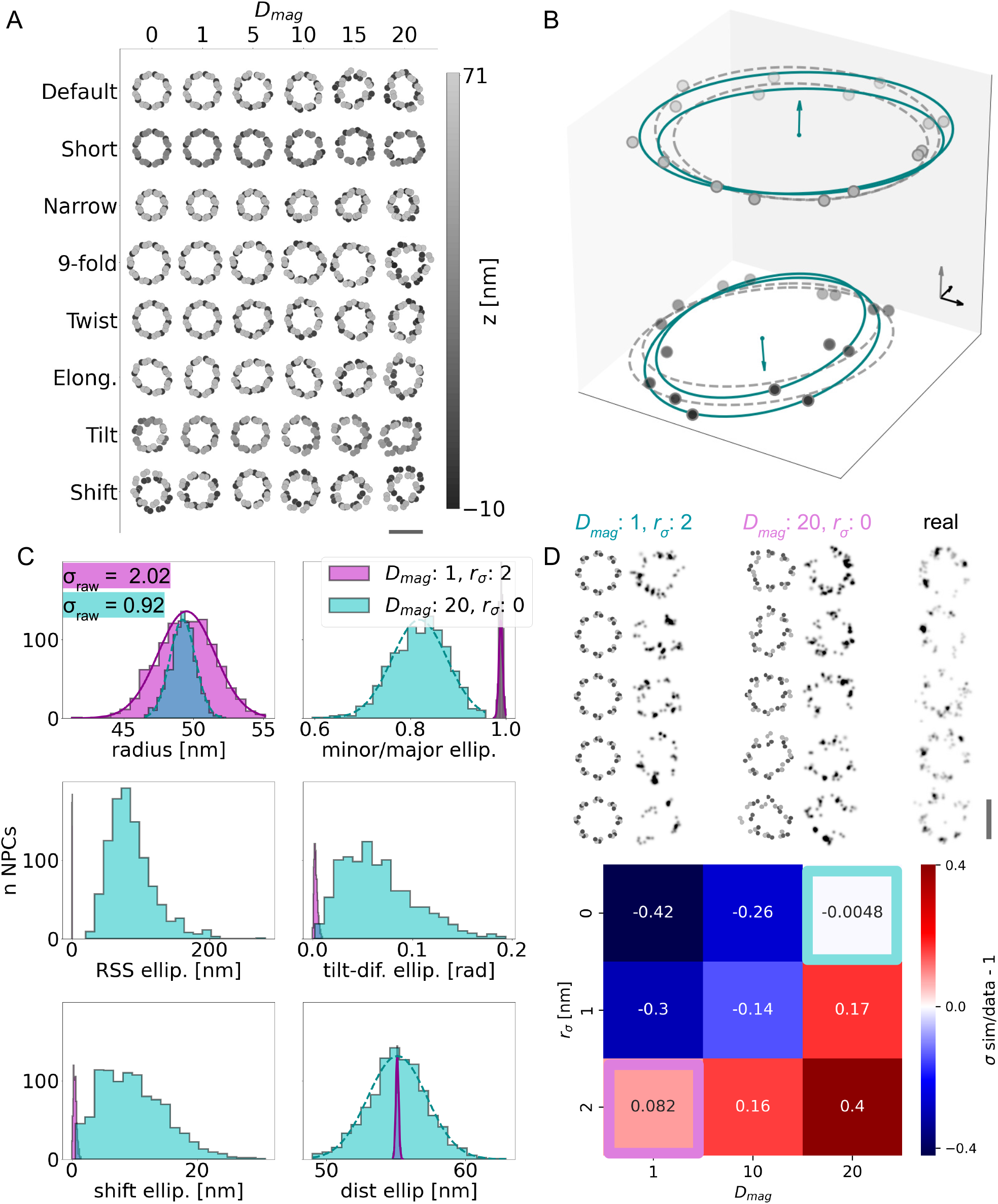
**A:** Different types of geometric deformation of Nup107 at increasing *D*_mag_ (and *O*_mag_. Scale bar 100 nm. **B:** Nup96 with *D*_mag_ = 20. Ellipse (continuous teal line) and circle (gray dashed line) fitted to each ring. Teal arrows indicate tilt of ORs (Suppl. section 9.2). Gray arrow indicates z direction. All arrows are 10 nm long. **C:** Distribution of extracted features of 1000 Nup96 with *D*_mag_ = 20 (magenta); and of 1000 Nup96 with *D*_mag_ = 1 and standard deviation of radius (*r*_*σ*_) = 2 nm (teal). The radius for an individual ground truth NPCs is determined by fitting a 3D circle to each of *g* rings and averaging *g* radii. Minor/major axis ellip. and RSS ellip. is determined equivalently, but by fitting ellipses. (Suppl. section 8). All other features are computed according to Suppl. section 9. Standard deviation of radii in raw data (no photophysics) is 2.02 for *D*_mag_ = 1 and (*r*_*σ*_) = 2 nm and 0.92 for *D*_mag_ = 20. **D:** top: Example NPCs, Nup96, from left to right: *D*_mag_ = 1, (*r*_*σ*_) = 2 nm, raw coordinates and with photophysics. *D*_mag_ = 20, (*r*_*σ*_) = 0 nm, raw coordinates and with photophysics. Corresponding real NPCs from Thevathasan et al. (2019) Bottom: Comparison of radial variability between simulated data (with photophysics, 1000 NPC per combination of *r*_*σ*_) and *D*_mag_ and real data: *r*_*σ*_= 2.1 nm, from Thevathasan et al. (2019) Magenta and teal boxes highlight example parameters shown in C and D, top.

Optionally, *f*_*i*_ may be modified to include other types of deformations (see Elongation, Suppl. section 7.6). Lastly, *f*_*i*_ is transformed into a vector 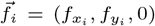 that is radial and planar to avoid rotations and flips with

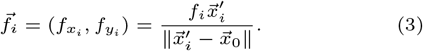

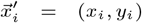 thereby is the x-y position of a node to which a 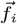 is applied. Any set of forces with no net-torque would avoid rotations, however radial forces are simpler to compute and more intuitive. Predominantly radial forces stemming from the NE are the most parsimonious explanation for observed dilations of the NPC, particularly as no motor activity of scaffold Nups has been observed (Zimmerli et al., 2021; Mosalaganti et al., 2022).

The final deformation of the model is computed for each ring separately (Suppl. section 4).

### Random axial offset is independent of springs

Axial resolution is poorer than lateral resolution in conventional SMLM (Huang et al., 2008). Deforming a spring-model in only two dimensions strongly speeds up computation and greatly simplifies the necessary spring layout. We therefore ignore spring interactions when offsetting nodes axially. Axial offset *o*_*i*_ is computed analogously to, but independently from, radial scalar forces (Eq. 1, 2, 3). The magnitude of axial offset *O*_mag_ is analogous to *D*_mag_ and set to *O*_mag_ = *D*_mag_/2 by default. The offset-vector on a node *i* is 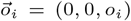. The updated node coordinates are therefore 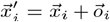. Following, *O*_mag_ is implied when specifying *D*_mag_.

### Simulating geometric variability

#### Geometric variability can be set as a fixed parameter or randomised

Geometric variability of the NPC is commonly observed in imaging data (Hinshaw and Milligan, 2003; Löschberger et al., 2014; Stanley, Fassati, and Hoogenboom, 2018; Beck et al., 2007; Maimon et al., 2012; Huijben et al., 2021; Heydarian et al., 2021; Stanley, Fassati, and Hoogenboom, 2018; Sabinina et al., 2021; Zimmerli et al., 2021; Mosalaganti et al., 2022; Sabinina et al., 2021; Heydarian et al., 2021; Curd et al., 2020). We simulate variability in radius, elongation, distance and twist angle between the nucleoplasmic- and cytoplasmic side, tilt and shift between the nucleoplasmic- and cytoplasmic side, rotational symmetry, and elongations (See Figure 4A for an overview and Table 1). Either of these variables *X* can be kept as standard input value, or redefined by the user. A measure of variation can be user-input for any parameter, except for the discrete rotational symmetry. This measure is the standard deviation *σ*if not indicated otherwise. If a standard deviation is given, a new value *X*^*′*^ for each model NPC is drawn from a normal distribution following

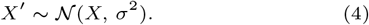

**Table 1.** Default and modified geometric parameters of simulated Nup107 (N terminus)

| Parameter | Symbol | Default | Modified value | Justification |
| --- | --- | --- | --- | --- |
| Mean radius | $\bar{r}$ | 48 nm | 38.4 (80% of 48 nm) | Observed<sup>1</sup> |
| Mean dist NR to CR | $\bar{d}$ | 58.6 nm | 30 nm | To resemble Nup133, to test z-resolution<sup>1</sup> |
| Elongation (short/long axis) | $q$ | 1 | 0.835 (0.8 at NPC) | Commonly observed<sup>2</sup> |
| Twist angle | $\theta$ | 9.6° | 14° | Observed<sup>3</sup> |
| Symmetry | $s$ | 8-fold | 9-fold | Commonly observed<sup>4</sup> |
| CR and NR tilt | $\kappa$ | None | 100 | Hypothetical |
| CR and NR shift | shift | None | $\sigma = 2$ nm | Hypothetical |

<sup>1</sup> Sabinina et al., 2021<sup>2</sup> Beck et al., 2007; Maimon et al., 2012; Huijben et al., 2021<sup>3</sup> Heydarian et al., 2021<sup>4</sup> Hinshaw and Milligan, 2003; Löschberger et al., 2014; Stanley, Fassati, and Hoogenboom, 2018

### Nups are assigned to Cytoplasmic and Nucleoplasmic side to simulate axial variability

The cytoplasmic side (CS) is defined as all *n* selected Nups *P*_*i*_, *i* ∈ 1 … *n*, with axial coordinate 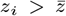, where 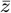 is the arithmetic mean of all *n* axial coordinates. A Nup is assigned to the nucleoplasmic side (NS) if 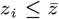.

#### Ring Distance

*d* is the difference between the average z positions of the CS 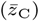 and NS 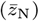 (Figure 4A “Short”, Suppl. section 7.2).

#### Tilt

the CS and NS can be tilted independently from each other. A tilt-vector for each side is sampled from a von Mises-Fisher distribution, which is a Gaussian distribution projected on a sphere. The NS and CS are then rotated to be orthogonal to their respective tilt-vectors. The mean direction of tilt-vectors is vertical, which is equivalent to no tilt. The dispersion of the von-Mises Fisher distribution is defined by user-input variable *κ*, where smaller values denote a higher dispersion, therefore more tilt (Figure 4A “Tilt”, Suppl. section 7.3).

#### Shift

the CS and NS side can be shifted laterally, representing shear. For each side, offset in x and y-direction are independently sampled from a Gaussian distribution with mean 0 and user-defined standard-deviation (Figure 4A “Shift”).

#### Twist Angle

we define the twist angle *θ* as the difference between the mean CS angles and the mean NS angles within one spoke, assuming a spoke contains the most acute possible *θ* (Figure 2, Figure 4A “Twist”, Suppl. section 7.4).

### Lateral variability

#### Radius

the arithmetic mean horizontal distance 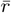 of Nups to the central axis. Changing the radius isotropically scales the NPC in lateral direction. Nups are therefore further apart in dilated NPCs (Figure 4A “Narrow”, Suppl. section 7.5).

#### Elongation

elongated NPCs are generated through approximation with an ellipse, whose ratio of short to long axis *q* is defined by the user. The rotational alignment between the NPC and this ellipse is randomised. Elongating forces 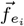 on nodes *i* ∈ 1 … *m* are finally sampled based on the ellipse and added to any other deforming forces (Figure 4A “Elong.”, Suppl. section 7.6).

#### Symmetry

for NPCs of varying symmetries, we keep the geodesic distance between Nups constant and preserve the absolute difference of radii between rings, as we assume that the space taken up by each Nup does not change with its copy number. Consistently, NPCs with 9-or 10-fold rotational symmetry exhibit a greater diameter (Hinshaw and Milligan, 2003; Löschberger et al., 2014; Stanley, Fassati, and Hoogenboom, 2018). Unlike for varying radii, the angle between Nups changes (Figure 4A “9-fold”, Suppl. section 7.7).

### Interplay between Geometric and Irregular variability

Irregular forces and correlated axial offset will deform the NPC model from its input geometric parameters, such as ring-distance, tilt, shift, radius and elongation (Figure 4C). Irregular variability can thus be disentangled into a geometric component and true irregularity, which represents irregular shapes visible in imaging data (Stanley, Fassati, and Hoogenboom, 2018; Beton et al., 2021; Löschberger et al., 2014). Our framework hence allows to tune the proportion of true irregularity, as well as the relative proportions of different types of geometric variability.

### Extracting features

Features are computed using parameters of fitted structures: we fit a 3D circle and 3D ellipse to each sub-complex, or to each NPC ring *g*. We define sub-complexes to be the Cytoplasmic ring (CR), Inner ring (IR), Nucleoplasmic ring (NR), the bridge between CR and IR (BRCR), and the bridge between IR and NR (BRNR) (Table S3, Suppl. section 8, 9).

### Output files and image data simulation

For each simulation experiment, NPC coordinates are saved in a CSV file that can be loaded into photophysics simulation software (e.g. Ries, 2020). A metadata file with properties of simulated NPCs is generated, as well as a CSV files containing features of circles and ellipses fitted to subcomplexes (CR, BRCR, IR, BRNR, NR), a CSV file with circle- and ellipse features fitted to individual rings, and a CSV file containing circle- and ellipse features per NPC obtained by averaging *g* ring-features. (Suppl. section 10).

## Results

### Simulated Dataset

We used PDB model 5A9Q (Appen et al., 2015) as basis for simulations due to its high pseudo-atomic resolution. We selected N-terminally labelled Nup107 (short: Nup107) and C-terminally labelled Nup96 (short: Nup96), (Tables S4, S3), which are frequently studied (e.g. Sabinina et al., 2021; Thevathasan et al., 2019; Wu, Hoess, et al., 2022; Li et al., 2022). We used Nup107 for a general purpose dataset, and Nup96 to recapture the specific imaging experiment carried out in Thevathasan et al. (2019). A virtual nucleus with 1000 strongly deformed NPCs (Nup96) takes 25 -31 s to simulate and 20 s to export as a CSV file (including features and metadata) on a personal laptop (Ubuntu Linux 20.04, processor model: i7-8550U, Processor frequency: 1.80 GHz, RAM: 16 GB).

#### N-Nup107

For Nup107, we simulated eight classes representing commonly observed or hypothesized variability: unmodified reference NPCs, smaller radius, smaller CR-NR distance, elongated NPCs, 14° twist angle, 9-fold symmetry, shifted CR and NR, and independently tilted NR and CR (Table 1, Figure 4A). For each class, we simulated NPCs with deformation mag 0, 1, 5, 10, and 15; 1000 NPCs each, and generated output files according to Methods “Output files and Image simulation”.

#### Nup96-C

Thevathasan et al. (2019) published a simple geometric analysis of Nup96 radii based on well-characterised, good-quality imaging data. We simulated comparable ground-truth NPCs by sweeping two parameters that contribute to the variability of fitted NPC radii: The standard deviation of NPC radius *r*_*σ*_, and irregular variability *D*_mag_. *D*_mag_ contributes to geometric variability, such as ellipticity, ring-tilt, ring-distance, and ring-shift, as well as to true irregular variability, measured as the RSS from ring-wise, 3D-fitted ellipses (Figure 4C, Suppl. section 9). The standard-deviation of radii *r*_*σ*_fitted to 2D NPCs in real-data was 2.1 nm (Thevathasan et al., 2019). We therefore simulate each 1000 NPCs with *r*_*σ*,model_ of 0, 1, and 2 nm; as well as *D*_mag_ of 1, 10, and 20, resulting in 9 combinations of *r*_*s*,model_ and *D*_mag_ (Figure 4D).

### Simulated Image Acquisition

Photophysics were simulated in SMAP (Ries, 2020). Photophysics parameters were modified from Wu, Hoess, et al. (2022), which represents a good-quality imaging experiment. Wu, Hoess, et al. (2022) simulate irregularity as a random, independent offset from the expected localisation position called “free linkage error”. Because our model accounts for shape irregularity with *D*_mag_, we are able to reduce the free linkage error to what it is expected to be in reality, namely the size of the fluorescent label (Yao et al., 2021). For Nup107, Simulation parameters are shown in Table S5. For Nup96, we further modify photophysics-parameters to correspond to Thevathasan et al. (2019) (Table S6). Simulated imaging data were exported as CSV files.

### Comparison with real NPCs

In our pipeline, parameters of simulated geometric-, and irregular shape variability can be adjusted, and features of resulting NPC epitopes can be directly read-out. This allows to systematically disentangle factors contributing to NPC variability. In combination with photophysics-simulations, this offers a reference to variability in real-data. We ran the radius-analysis from Thevathasan et al. (2019) on our simulations and compared *r*_*σ*_of real and simulated data with

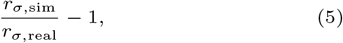

such that negative values indicate too little simulated variability, and positive value too much simulated variability.

The radius-variability of real NPCs can both be achieved by predominantly varying the underlying model-radius, as well as by only simulating irregular variability (Figure 4C, D). Assuming that the simulated photo-physics accurately capture real-life photophysics, we can exclude combinations of *r*_*σ*,model_ and *D*_mag_ that have too much or too little radial variability.

## Discussion

Current SMLM simulation methods cover the entire imaging pipeline (Sinkó et al., 2014; Venkataramani et al., 2016; Novák et al., 2017; Griffié et al., 2020; Sage et al., 2019; Ries, 2020) including basic variations of NPC geometry (Wu, Hoess, et al., 2022; Huijben et al., 2021). However, no approaches allow simulating irregular but coherent shape variations. Here, we introduce CIR4MICS, a toolbox that simulates irregularly shaped NPCs as well as a wide range of geometric NPC variability reported in literature and anecdotally. We further provide a reference dataset that includes deformed ground truth NPCs, annotated features, as well as corresponding simulated imaging data using SMAP (Ries, 2020). Amongst other applications, our software allows to compare fluorescence microscopy clustering, classification, and averaging methods. A side-by-side comparison with real data allows to find simulation parameters that best represent real NPCs, thus elucidating their structural variability.

We integrated pseudo-atomic models of human NPCs in membrane isolates (Appen et al., 2015; Kosinski et al., 2016; Mosalaganti et al., 2022) and in their native environment (Schuller et al., 2021; Mosalaganti et al., 2022) into CIR4MICS. We additionally provide information and scripts on how to integrate further models. Simulations can therefore be extended to NPCs of other species such as yeast (Allegretti et al., 2020; Kim et al., 2018; Zimmerli et al., 2021), especially since advances in integrative modelling (Rantos, Karius, and Kosinski, 2021; Mosalaganti et al., 2022) promise a wider gamut of resolved NPCs. Generally, CIR4MICS allows to integrate any rotationally symmetric structure, such as cilial cross-sections, with little to no modifications (Shi et al., 2019). Filamentous structures could further be simulated by increasing the model symmetry such that local topology resembles a line segment. Lastly, simulated epitope number and distance can be freely adjusted, providing a reference for any application where structure does not matter.

In our framework, Nup coordinates are transformed into a spring model to which static deforming forces are applied to achieve irregular, yet coherent shapes. Our simulations resemble coarse grained elastic models (Wolf and Mofrad, 2008; Lezon, Sali, and Bahar, 2009) or molecular dynamics simulations (Mosalaganti et al., 2022) in that a model of the NPC is subjected to deforming forces. Crucially however we do not intend to predict true NPC structural variability, but to replicate observed or presumed variability accompanied by synthetic ground truth. Such hypothesis-driven approaches benefit from fast computations as they require multi-parameter sweeps, as well as a rapid turnover of updated hypotheses and simulations. Accordingly, our simple model layout allows to generate a virtual nucleus of 1000 annotated NPCs in under one minute on a standard laptop.

Our software allows to examine how a wide range of image analysis methods behave on simulations, and to directly compare the simulated output to real data. Clustering and classification methods can be studied based on their behaviour on real underlying classes, such as distinct geometries (Theiss et al., 2022), as well as on increasing levels of shape irregularity. Particle averaging methods align degraded observations to reconstruct their underlying structure, but struggle with biological heterogeneity, which our method can simulate. Notably, a method that evens out deformed circular structures before averaging has been developed (Shi et al., 2019) and could be tested on simulated NPC shape irregularity. Furthermore, relative distances of localisations of multiple structures can be statistically analysed to extract information of the underlying average geometry (Curd et al., 2020). CIR4MICS allows to study the effect of biological variability on such analyses.

Geometric model fitting allows to characterise the underlying structure in terms of geometric parameters, which can form the basis for clustering, classification, and particle averaging (Wu, Hoess, et al., 2022; Thevathasan et al., 2019). Mismatch between a geometric model and the underlying structure will affect accuracy and precision of fitted parameters. (Wu, Hoess, et al., 2022). We elaborate on this notion of mismatch by simulating biological shape irregularity as parameter *D*_mag_. *D*_mag_ contributes to geometric variability, such as ellipticity, ring-tilt, ring-distance, and ring-shift, as well as to true irregularity, such as the RSS from an optimally fitting geometric representation. By adjusting *D*_mag_ and geometric parameters, we can finetune the proportions of distinct types of variability in a dataset. Reproducing a simple circle-fitting geometric analysis (Thevathasan et al., 2019) on simulated data, we showed that both predominantly geometric variability and predominantly irregular variability can replicate variability of radii found in real data. Previous simulations do not allow to distinguish between true radial variability and coherent shape irregularity (Wu, Hoess, et al., 2022; Venkataramani et al., 2016; Huijben et al., 2021). Assuming equal imaging noise, we can conclude that any combination of biological variability parameters that exceeds or undershoots variability of real data, does not represent real NPCs. Here, we determined imaging noise settings through a literature search (Wu, Hoess, et al., 2022; Thevathasan et al., 2019; Yao et al., 2021). Simulated imaging noise could however be further adjusted via a side-by-side comparison of real and simulated image quality. Additionally, a similar comparison of real and simulated NPCs could be rerun with exceptional-quality data (Diekmann et al., 2020), and more sophisticated geometric data analyses (Wu, Hoess, et al., 2022; Sabinina et al., 2021). Comparing a wider range of geometric properties, will also allow to narrow down the possible geometric variations of NPCs while refining simulation priors.

The analysis we present here assumes an unimodal distribution of NPC geometry. It is uncertain whether geometric parameters of real NPCs follow a unimodal or multi-modal distribution. For example, NPCs are commonly eight-fold symmetric, however nine- and ten-fold symmetric NPCs have been reported (Hinshaw and Milligan, 2003; Löschberger et al., 2014; Stanley, Fassati, and Hoogenboom, 2018). Other aspects of NPC geometry might be truly uni-modal; or multi-modal, but not detectable by analysis software; or uni-modal with clustering methods falsely reporting multi-modality. Crucially, CIR4MICS can generate synthetic ground truth that would allow to establish the limits of SMLM data and analysis methods to discriminate between different NPC geometries. Simulations allow to assess the amount and quality of data needed to answer a particular question. Synthetic data hence aids in the choice of analysis methods, helps develop new methods, and informs experimentalists about the feasibility and requirements of a given analysis. Users can for instance use the a pre-genrated two-class data-set, say NPCs with twist-angle of 9.6° and 14°(Heydarian et al., 2021), to test whether an analysis method recovers this bi-modality. Results can be compare to equivalent analysis on real-NPCs. Users can further sweep parameters to test the breaking points when bi-modality is no longer recoverable: The difference in angle can be increased or reduced and angles can be drawn from a Gaussian distribution with increasing standard deviation. The shape irregularity *D*_mag_ can be increased and the number of simulated NPCs per class can be adjusted to find the minimum number required for statistical significance. A uni-modal distribution of angles with increasing standard-deviation should be analysed for comparison.

CIR4MICS combines microscopy simulations with molecular physics simulations. Notably, it is not limited to SMLM simulations, as several other microscopy techniques work on the basis of attaching fluorescent or electron-dense labels to a structure, the locations of which can be expressed by coordinates. Such coordinates generated by our node and spring model can therefore be passed to simulation pipelines of other microscopy techniques. More such combinations of simulation or modelling techniques could emerge in the future, for instance the integration of Brownian dynamics or molecular dynamics with any kind of imaging simulations. Such interfaces would result in more realistic image simulations, and allow to directly compare dynamics simulations with real data.

## Code and Data availability

All described code is available at https://github.com/uhlmanngroup/cir4mics. Simulated data is available at BioStudies (Accession number S-BSST1058).

## Acknowledgements

We thank Jeremy Tempkin, Andrej Šali, Wanlu Zhang, Vilma Jimenez Sabinina, Zhichao Miao, Gerard Kleywegt, and Jan Kosinski for discussions. Thanks to Ben Woodhams for testing our software. Special thanks to Yu-Le Wu and Philipp Hoess for discussions and help with SMAP. This work was supported by Wellcome Trust [grant number 212962/Z/18/Z to C.R.].

