## Supplementary material for "Simulating structurally variable Nuclear Pore Complexes for Microscopy"

### 1 Notation

Information on notation can be found in Table S1.

**Table S1.** Important notation

| Name | Variable | Note |
|---|---|---|
| Symmetry | $s$ | Default $s = 8$ |
| Copy number of Nup, mass of node | $m$ | |
| Number of rings | $g$ | $m = gs$. Not the same as CR, NR, IR |
| Coordinates of node $i$ | $\vec{x}_i = (x_i, y_i, z_i)$ | |
| Central anchor coordinates | $\vec{x}_0$ | At (0, 0) |
| Centre coordinates | $\vec{x}_c$ | Not necessarily at (0,0) |
| Indices | $i, j, l$ | |
| Radius | $r$ | |
| Nup of interest $i$ | $nup_i$ | |
| Individual Nup/point | $p$ | |
| Type of Nup | P | |
| Cytoplasmic or Nucleoplasmic side | C, P | |
| Number of connected neighbours of a node | $2h$ | |
| Spring constant | $k$ | |
| Magnitude of lateral deformation | $D_{\text{mag}}$ | |
| Magnitude of axial offset | $O_{\text{mag}}$ | |
| Standard deviation, RBF kernel free parameter | $\sigma$ | |
| Arithmetic mean of x | $\bar{x}$ | |
| Ring distance, damping constant | $d$ | |
| Angle of Nup $i$ within RU | $\alpha_i$ | |
| Azimuthal angle between CR and NR | $\theta$ | Also called “twist angle” |
| Argument of parametric form of of ellipse and evolute, | $\tau$ | (58), (59) |
| Rotational angle of ellipse | $\rho$ | |
| Rotation whole NPC | $\delta$ | |
| Generic max count | $n$ | |
| Semiminor axis ellipse | $b$ | |
| Semimajor axis ellipse, acceleration | $a$ | |
| Semiminor/semimajor axis ratio ellipse | $q$ | $q = b/a$ |
| Circumference | $c$ | |
| Scale factor,<br>Polynomial coefficient in implicit form of ellipse | $C$ | |
| Sign | sgn | $+, -, 0$ |
| Geodesic distance of a circle segment | $d_{\text{segment}}$ | segment described as angle |
| Euclidean distance between $\vec{x}_i$ and $\vec{x}_j$ | $\|\vec{x}_i - \vec{x}_j\|$ | |
| Cross product of $\vec{x}_i$ and $\vec{x}_j$ | $\vec{x}_i \times \vec{x}_j$ | |
| Dot product of $\vec{x}_i$ and $\vec{x}_j$ | $\vec{x}_i \cdot \vec{x}_j$ | |
| Magnitude of $\vec{x}$ | $\|\|\vec{x}\|\|$ | |

#### 1.1 NPC as pointcloud

The NPC or any substructure of it can be represented as point-cloud $X \in \mathbb{R}^{m \times 3}$, where $m$ is the total number of points (=nodes) in $X$ and 3 is the number of spatial dimensions:

$$X = \begin{bmatrix} x_1 & y_1 & z_1 \\ \vdots & \vdots & \vdots \\ x_i & y_i & z_i \\ \vdots & \vdots & \vdots \\ x_m & y_m & z_m \end{bmatrix} \tag{1}$$

#### 1.2 Rodrigues rotation

Point clouds (e.g. (1)) or vectors can be rotated using Rodrigues rotation

$$X_{\text{rot}} = X\cos(\theta) + (\hat{k} \times X)\sin(\theta) + \hat{k}(\hat{k} \cdot X)(1 - \cos(\theta)) \tag{2}$$

where $\hat{k}$ is the axis of rotation and $\theta$ is the angle of rotation.
This can be compactly expressed as:

$$X_{\text{rot}} = R(\hat{k}, \theta)X \tag{3}$$

### 2 Extracting Nup Coordinates from PDB

#### 2.1 Finding the model centre

The model centre is equivalent to the axis of rotational symmetry. We find the model centre $\vec{x}_c$ by calculating the midpoint of any Nup $i$ with coordinates $\vec{x}_i = (x_i, y_i)$, $i \in 1 \dots s$, and the opposite Nup of the same type and on the same ring with index $j = \text{mod}(i + 0.5s, s)$ (4), provided $s$ is even. We verify $\vec{x}_c$ for multiple opposing Nup pairs. For models 7PEQ, 7PER [1], 7R5K, 7R5J [2], and 5A9Q [3], $\vec{x}_c$ corresponds to the midpoint of the underlying electron microscopy maps.

$$\vec{x}_c = 0.5(\vec{x}_i + \vec{x}_j) \tag{4}$$

#### 2.2 Finding radii and angles of Nups within one Rotational unit

Nup $\vec{p}_i$ with $i = 1$, $i \in 1 \dots n$, is set as reference $\vec{p}_{\text{ref}} = (x_{\text{ref}}, y_{\text{ref}})$, where $n$ is the total number of individual Nups within a rotational unit (RU) selected for labelling regardless of color channel. Here, the axial dimension can be omitted due to the underlying model's planarity. $\alpha_i$ of $\vec{p}_i = (x_i, y_i)$ relative to $\vec{x}_c$ and $\vec{p}_{\text{ref}}$ is calculated via (5). Angles are measured in radians if not indicated otherwise and anticlockwise angles are positive.

$$\alpha_i = \text{atan2}(y_i - y_c, x_i - x_i) - \text{atan2}(y_{\text{ref}} - y_c, x_{\text{ref}} - x_c) \tag{5}$$

If $\alpha_i > \pi$ or if $\alpha_i < -\pi$ it is represented as the explementary angle $\alpha_i - 2\pi$ or $\alpha_i + 2\pi$ respectively. The mean angle $\bar{\alpha}$ is computed as

$$\bar{\alpha} = \sum_{i=1}^{n} \frac{\alpha_i}{n}, \tag{6}$$

and used to zero-centre $\alpha_i$ following

$$\alpha'_i = \alpha_i - \bar{\alpha} \text{ for } i \in 1 \dots n. \tag{7}$$

For multi-channel-simulations, the first type of Nup, $P_1$, can be optionally set as a reference Nup $P_{\text{ref}}$. $P_{\text{ref}}$ contains $o$ individual Nups $p_j$ in one RU with corresponding $\alpha_j$, $\bar{\alpha}_{\text{ref}}$ is computed via

$$\bar{\alpha}_{\text{ref}} = \sum_{j=1}^{o} \frac{\alpha_j}{o}, \tag{8}$$

and $\alpha_i$ is updated such that $\bar{\alpha}_{\text{ref}} = 0$ with

$$\alpha''_i = \alpha'_i - \bar{\alpha}_{\text{ref}} \text{ for } i \in 1 \dots n. \tag{9}$$

Lastly, $r_i$ is the Euclidean distance between coordinates $\vec{x}_i$ of $p_i$ and $\vec{x}_c$, defined as

$$r_i = \|\vec{x}_i - \vec{x}_c\|\,, \tag{10}$$

and is converted to nm.

### 3 Layout of the spring model

Spring models ensure that any offset on nodes distributes throughout the entire model, as represented proteins are directly or indirectly connected to their neighbours (Figure S1A). Nodes are connected to their $2h$, $h \in 0 \ldots \lfloor s/2 \rfloor$, neighbours via circumferential springs. Higher values of $h$ require higher deforming forces to achieve sufficiently irregular deformations. Computation time would therefore increase due to these higher forces, and due to the additional number of springs. By contrast, small values for $h$ lead to overly jagged deformations. $h$ can be set by the user and is 2 by default. Furthermore, the spring constant $k$ of the shortest circumferential spring is 1 and decreases inversely proportional with spring length. If $h$ is set to $\lfloor s/2 \rfloor$, the nodes are fully connected. Consequently, if $s$ is even, opposite nodes are connected by two springs. $k$ for these springs are therefore halved to make them equivalent to a single spring. Furthermore, spring constant $k$ of radial springs is 0.7, a value lower than that of the shortest circumferential spring, as we assume protein-protein interactions to be stronger than protein-membrane interactions. A lower value would lead to more drift upon deformation and concomitantly to longer computation times.

### 4 Ordinary differential equations compute deformation

Deformations are computed for each ring separately. Nodes are therefore $i \in 1 \ldots s$. To calculate the position $\vec{x}_i = (x_i, y_i, z_i)$ of a node $i$ in response to a force $\vec{f}_i = (f_{x_i}, f_{y_i}, 0)$, Newton's second law of motion can be consulted:

$$\vec{f}_i = m_i \vec{a}_i. \tag{11}$$

Henceforth, we assume the mass of the node $m_i = 1$. The acceleration of the node $\vec{a}_i$ is the second derivative of $\vec{x}_i$ over time $t$, which changes (11) into

$$\frac{d^2\vec{x}_i}{dt^2} = \vec{f}_i. \tag{12}$$

This only holds true for a disconnected node. Hooke's spring law states that a force stored in a spring between nodes $i$ and $j$ depends on spring constant $k_{ij}$, and the displacement of the spring from its resting length $l_{ij}$. In a spring system with $s$ nodes, the force $\vec{f}_{is}$ from the system on a node $i$ is the sum of the forces stored in each spring, given by

$$\vec{f}_{is} = \sum_{j=0, j\neq i}^{s} k_{ij} \frac{\vec{x}_i - \vec{x}_j}{\|\vec{x}_i - \vec{x}_j\|} \left(l_{ij} - \|\vec{x}_i - \vec{x}_j\|\right). \tag{13}$$

Here, $j = 0$ for an anchor node at $\vec{x}_0$ and $j \in 1 \ldots s$ for the NPC nodes.
The acceleration, and therefore position $\vec{x}_i$ of a node $i$ over time thus depends on both an external force $\vec{f}_i$, and the force $\vec{f}_{is}$ stored in the system according to

$$\frac{d^2\vec{x}_i}{dt^2} = \vec{f}_i + \vec{f}_{is}. \tag{14}$$

Due to the conservation of energy, displacing a node from its equilibrium would result in continuous movements of the node. Damping $-d\frac{d\vec{x}_i}{dt}$ needs to be applied to the node, so that it reaches a new equilibrium given by

$$\frac{d^2\vec{x}_i}{dt^2} = \vec{f}_i + \sum_{j=0, j\neq i}^{s} k_{ij} \left( \frac{\vec{x}_i - \vec{x}_j}{\|\vec{x}_i - \vec{x}_j\|} (l_{ij} - \|\vec{x}_i - \vec{x}_j\|) \right) - d\frac{d\vec{x}_i}{dt}, \tag{15}$$

where $d$ is a damping constant and $\frac{d\vec{x}_i}{dt}$ is the velocity of the node.
We are interested in static variability. To obtain solutions at equilibrium, we numerically solve the system using scipy.integrate.solve_ivp [4] until time $t$, where $t$ is large.

### 5 Forces are correlated by distance

Covariance between forces on nodes $\vec{x}_i = (x_i, y_i, z_i)$ and $\vec{x}_j = (x_j, y_j, z_j)$, with $i, j \in 1 \ldots m$ are given by the RBF kernel,

$$\text{cov}(\vec{x}_i, \vec{x}_j) = e^{-\frac{\|\vec{x}_i - \vec{x}_j\|^2}{2\sigma^2}}, \tag{16}$$

to preserve local structures of nearby nodes while ensuring irregularity.
Here, $\sigma$ is a free parameter. We couple $\sigma$ to the size of the model by setting it to a fraction of the mean radius $\bar{r}$ (Figure S1B). We set $\sigma = \bar{r}/2$ as default, as it was empirically observed to produce the best results.

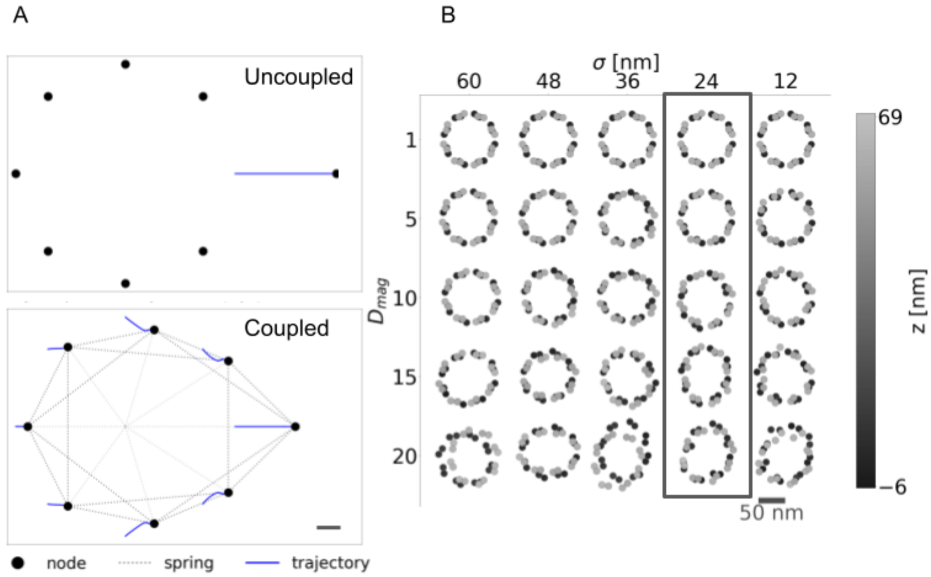


**Figure S1. A:** Behaviour of a spring model upon application of a point force. Purely geometric models are incapable of propagating a point force, whereas a spring model allows for irregular shape variations. Scale bar 10 nm **B:** RBF $\sigma$ vs deformation mag. Large values of $\sigma$ prohibit non-convex deformation, while deformations are too irregular for small values of $\sigma$. We set $\sigma$ to $\bar{r}/2$ by default, which is 24 nm in this case. Boxed are reasonable values for $\sigma$ and deformation mag. Scale bar 100 nm, Nup107 N-terminus of 5A9Q. Color-bar indicates z-position in nm. [3].

### 6 Dynamics simulations for non-NPC structures

As temporal and spatial resolution of SMLM counter-correlate, it is near impossible to resolve NPC shape and dynamics. [5]. Model-properties such as symmetry, spring-damping, and correlation of deforming forces can however be adjusted to achieve membrane-like or filament-like structures with pseudo-chaotic dynamics. As such dynamics are challenging, or impossible, to predict, our simulations could benchmark or train dynamics reconstruction software for SMLM [6], [7].

### 7 Geometric variability

#### 7.1 Assigning Nups to Cytoplasmic and Nucleoplasmic side

To assign Nups to the cytoplasmic side (CS) or nucleoplasmic side (NS), we compute the average z position $\bar{z}$ of all selected Nups. Rings $i$ with $i \in 1 \ldots g$ are assigned to the CS, if $z_i > \bar{z}$ and to the NS if $z_i \leq \bar{z}$.

#### 7.2 Ring distance

the distance $d$ is the difference between the average z positions of Nups on the CS, $\bar{z}_{\text{CS}}$ and the NS, $\bar{z}_{\text{NS}}$.

$$d = \bar{z}_{\text{CS}} - \bar{z}_{\text{NS}} \tag{17}$$

When a new distance $d'$ is set manually or randomly sampled, the axial coordinates $\bar{z}_{\text{CS}}$ are updated to $\bar{z}'_{\text{CS}}$.

$$\bar{z}'_{\text{CS}} = \bar{z}_{\text{CS}} - (d - d') \tag{18}$$

#### 7.3 Tilt

##### 7.3.1 Sampling tilt vectors

The code was adapted from `https://dlwhittenbury.github.io/ds-2-sampling-and-visualising-the-von-mises-fisher-distribution-in-p-dimensions.html`. A tilt-vector $\vec{x} = (x_x, x_y, x_z)$ is sampled from a von Mises-Fisher distribution, which is a Gaussian distribution projected on a sphere with mean orientation $\vec{\mu} = (\mu_x, \mu_y, \mu_z)$.
Initially, a vector $\vec{x}' = (x'_x, x'_y, x'_z)$ is sampled with a mean orientation $\vec{\mu}' = (1, 0, 0)$ and ultimately rotated to $\vec{x}$ with $\vec{\mu}$. In our case, $\vec{\mu} = (0, 0, 1)$ to reflect a planar orientation of the NPCs.
The vector $\vec{x}'$ can be computed via tangent-normal decomposition with a normal component $t\vec{\mu}'$ and a tangential component $\sqrt{1-t^2}\vec{\xi}$ [8], [9] following

$$\vec{x}' = t\vec{\mu}' + \sqrt{1-t^2}\vec{\xi}. \tag{19}$$

The value of $\vec{\xi} = (0, \xi_y, \xi_z)$ is uniformly sampled from a unit-circle [10] according to

$$\begin{aligned} \vec{\xi}'' &= (\xi''_y, \xi''_z) \ \sim \mathcal{N}(0,1)^2, \\ \vec{\xi}' &= \frac{\vec{\xi}''}{||\vec{\xi}''||}, \\ \vec{\xi} &= (0, \vec{\xi}'). \end{aligned} \tag{20}$$

Next, $t$ is sampled from the marginal of the von-Mises Fisher distribution in the interval $[-1, 1]$ corresponding to the upper and lower bounds in normal direction on the unit-sphere. It is obtained by rejection sampling [8], [9] following

$$b = \frac{2}{2\kappa + \sqrt{4\kappa + 4}}, \tag{21}$$

$$x_0 = \frac{1-b}{1+b}, \tag{22}$$

$$c = \kappa x_0 + 2\ln(1 - x_0^2), \tag{23}$$

whereby $\kappa$ is the concentration of the distribution.

- The values $Z$ and $U$ are sampled to compute $W$ as follows:

$$Z \sim \text{Beta}\,(1,1)\,, \tag{24}$$

$$U \sim \text{Uniform}(0,1), \tag{25}$$

$$W = \frac{1-(1+b)Z}{1-(1-b)Z}. \tag{26}$$

- If $t = W \Rightarrow \kappa W + (p-1)\ln(1 - x_0 W) - c \geq \ln(U)$ does not hold true, the process is repeated from (24).

Next, $\vec{x}'$ is computed via (19).
To rotate $\vec{x}'$ to $\vec{\mu}$, an orthonormal basis $O$ for the null-space of $\vec{\mu}$ is constructed. $\vec{\mu}^T$ and $O$ are concatenated to the rotation matrix $R$ given by

$$R = \begin{bmatrix}[\mu^T] & O\end{bmatrix}. \tag{27}$$

Finally, $R$ is applied to $\vec{x}'$ to obtain $\vec{x}$ as

$$[\vec{x}] = R[\vec{x}'^T]. \tag{28}$$

###### 7.3.2 Applying tilt

The NPC substructure to be tilted is represented as a point-cloud $X \in \mathbb{R}^{m\times 3}$, where $m$ is the total number of points (=nodes) in $X$ and 3 is the number of spatial dimensions (1).
$X$ is 0-centred in z-direction by subtracting the arithmetic mean $\bar{z}$ of $z_1 \ldots z_m$ following

$$X_{z0}(:,3) = X(:,3) - \bar{z}. \tag{29}$$

$X_{z0}$ are finally rotated to be orthogonal to the tilt vector $\vec{x}$ using Rodrigues rotation (3) as

$$X_{r,z0} = R(\hat{k},\theta)X_{z0}. \tag{30}$$

The axis of rotation is given as $\hat{k} = (\hat{x} \times \hat{e}_z^T)/||\hat{x} \times \hat{e}_z^T||$, and $\theta = \arccos(\hat{x} \cdot \hat{e}_z)$ is the angle of rotation, where $\hat{x}$ is the normalised tilt vector and $\hat{e}_z = (0,0,1)$ is the basis-vector in z.
Finally, the z-offset on $X_{r,z0}$ is reverted to obtain the final tilted point-cloud $X_r$,

$$X_r(:,3) = X_{r,z0}(:,3) + \bar{z}. \tag{31}$$

##### 7.4 Twist Angle

Selected Nups are assigned to the CS or NS. If $n > 1$ rings of the NoI occur on the CS, the arithmetic mean $\bar{\alpha}_{\mathrm{C}}$ of the ring angles defined as

$$\bar{\alpha}_{\mathrm{C}} = \sum_{i=1}^{n} \frac{\alpha_{i,\mathrm{C}}}{n}, \tag{32}$$

is used for further calculations. The value of $\bar{\alpha}_{\mathrm{N}}$ is computed analogously.
Furthermore, a RU does not always contain Nups with the most acute (closest to 0) angular distance $\theta = \bar{\alpha}_{\mathrm{N}} - \bar{\alpha}_{\mathrm{C}}$. Therefore, we probe the next clockwise- (cw) $\bar{\alpha}_{\mathrm{C,cw}} = \bar{\alpha} - 2\pi/8$ and anticlockwise (acw) angle $\bar{\alpha}_{\mathrm{C,acw}} = \bar{\alpha} + 2\pi/8$, and set the final twist angle $\theta$ to whichever $\bar{\alpha}_{\mathrm{N}} - \bar{\alpha}_{\mathrm{C,cw}}$, $\bar{\alpha}_{\mathrm{N}} - \bar{\alpha}_{\mathrm{C}}$, or $\bar{\alpha}_{\mathrm{N}} - \bar{\alpha}_{\mathrm{C,acw}}$ is the most acute.
This angle $\theta$ can then be changed to a user-defined $\theta'$ following

$$\begin{aligned} \alpha'_{i,\mathrm{C}} &= \alpha_{i,\mathrm{C}} + \frac{\theta - \theta'}{2}, \\ \alpha'_{i,\mathrm{N}} &= \alpha_{i,\mathrm{N}} - \frac{\theta - \theta'}{2}, \end{aligned} \tag{33}$$

##### 7.5 Radius

A new mean radius $\bar{r}'$ is set manually or randomly sampled. The radius of a Nup $p_i$ is denoted as $r_i$ with $i \in 1 \ldots n$, where $n$ is the total number of individual Nups in the simulated NPC. The $r'_i$ of individual Nups is updated following

$$r'_i = \frac{\bar{r}'}{\bar{r}}. \tag{34}$$

The angle between Nups is thereby kept constant.

##### 7.6 Elongation

Elongated NPCs are simulated by computing deforming forces from an ellipse with user-defined semiminor/semimajor axis ratio $q$. To this end, the circumference $c_j$ of each NPC ring $j \in 1 \ldots g$ is determined by $c_j = r_j 2\pi$. Next, semiminor- and semimajor axes $b_j$ and $a_j$ of an ellipse with circumference $\sim c_j$ are determined with

$$\begin{aligned} c_j &\approx 2\pi \sqrt{\frac{a_j^2 + b_j^2}{2}}, \\ a_j &\approx \frac{\frac{c_j \sqrt{2}}{2\pi}}{\sqrt{1 + q^2}}, \\ b_j &\approx q a_j. \end{aligned} \tag{35}$$

The resulting ellipse can be described with

$$\frac{x^2}{a_j^2} + \frac{y^2}{b_j^2} = 1, \tag{36}$$

where $x$ and $y$ are Cartesian coordinates.
NPC Cartesian coordinates are converted to polar coordinates and rotated by angle $\delta \sim U[-\pi, \pi]$ to vary the alignment of nodes with the later superimposed ellipse. Next, polar coordinates are converted back to Cartesian coordinates.

To project NPC coordinates onto the resulting ellipse, a scale factor $C_i$ needs to be found for each node $i \in 1 \dots s$ such that $C_i(x_i, y_i) = (x, y)$. The value of $C_i$ can be computed by substitution in (36) as

$$\begin{aligned} \frac{(C_i x_i)^2}{a_j^2} + \frac{(C_i y_i)^2}{b_j^2} &= 1, \\ C_i &= \sqrt{\frac{a_j^2 b_j^2}{x_i^2 b_j^2 + y_i^2 a_j^2}}. \end{aligned} \tag{37}$$

The difference $\triangle\vec{x}_i = (\triangle x_i, \triangle y_i)$ between the ellipse and NPC coordinates is computed as

$$\triangle\vec{x}_i = (\triangle x_i, \triangle y_i) = C_i(x_i, y_i) - (x_i, y_i). \tag{38}$$

Next, the compressive ($\text{sgn}(x) < 0$) or expanding ($\text{sgn}(x) > 0$) forces $\vec{f}_{e_i}$ on a node $i$ are determined with

$$\vec{f}_{e_i} = \text{sgn}(\triangle\vec{x}_i \cdot \vec{x}_i) \left\| \triangle\vec{x}_i \right\|. \tag{39}$$

The forces $\vec{f}_{e_i}$ therefore deform the NPC into an ellipse-like shape and are added to $\vec{f}_i$ from (3) in the main text.

##### 7.7 Symmetry

Computations are performed separately for each NPC ring. To determine the new radius $r'$ of an NPC with deviant symmetry $s$, a planar, isosceles triangle is constructed between two neighbouring Nups on the same ring, and the NPC central axis. The angle at the centre is $\beta = 2\pi/s$. The other two corners both have an angle of $\gamma = 0.5(\pi - \beta)$. Half the distance between neighbouring Nups $b_{\frac{1}{2}}$ is calculated via $b_{\frac{1}{2}} = r \sin(\pi/8)$, where $r$ is the radius of the 8-fold symmetric NPC ring. The updated radius $r'$ now corresponds to the hypotenuse of the right triangle between a Nup, the half-point to its neighbouring Nup, and the centre

$$r' = \frac{b_{\frac{1}{2}}}{\cos(\gamma)}. \tag{40}$$

If a Nup occurs in multiple rings $i \in 1 \dots g$, each with radii $r_i$, their initial ($\bar{r}$) and updated ($\bar{r'}$) arithmetic means are computed. Next, $r'_i = \triangle r_i + \bar{r'}$ with $\triangle r_i = r_i - \bar{r}$. Consequently, the radial distances between Nups in different rings are preserved.
For all rings, the angles $\alpha_i$ of Nups within one RU are updated to preserve the geodesic distance of adjacent Nups. First, the geodesic distance of Nups in an NPC with $s = 8$ is calculated via $d_{\alpha_i} = \alpha_i \bar{r}$. The updated angles $\alpha'_i$ are calculated via $\alpha'_i = d_{\alpha_i}/\bar{r'}$ (Figure 4A, 9-fold).

### 8 Fitting Circles or Ellipses to analyse deformed NPCs

##### 8.1 Non-planar NPCs

The code was adapted from `https://meshlogic.github.io/posts/jupyter/curve-fitting/fitting-a-circle-to-cluster-of-3d-points/`. We determine the planarity of point-clouds $X$ (1) representing the NPC or an NPC substructure with $X \in \mathbb{R}^{m \times 3}$, where $m$ is the total number of points (=nodes) in $X$. Individual points are denoted $\vec{x}_i = (x_i, y_i, z_i)$ with $i \in 1 \dots m$.
To this end, we determine the best-fitting plane such that the square sum of orthogonal distances of points to the plane is minimised. The centre of the point cloud is defined as the

arithmetic mean $\bar{x} = (x_c, y_c, z_c)$ of all points $\vec{x}_i$. The points $\vec{x}_i$ are 0-centred, $\vec{x}_{i,0} = x_i - \bar{x}$, resulting in a point-cloud $X_0$.
A singular value decomposition on $X_0$ yields $X_0 = U\Sigma V^T$, where the unit-vectors $V^T(1,:) = \hat{t}_1$ and $V^T(2,:) = \hat{t}_2$ are tangential to the best-fitting plane and point in the direction of the highest- and second-highest spread of $X_0$. The vector $V^T(3,:) = \hat{n}$ is the normal unit vector of the best-fitting plane.
Point clouds are planar if $\hat{n} = \hat{e}_z$, where $\hat{e}_z$ is the basis vector in z. If $\hat{n} \neq \hat{e}_z$, $X_0$ is projected onto the fitting plane using Rodrigues rotation (3) following

$$X_\text{r} = R(\hat{k}, \theta)X_0, \tag{41}$$

with $\hat{k} = (\hat{n} \times \hat{e}_z^T)/||\hat{n} \times \hat{e}_z^T||$ and $\theta = \arccos(\hat{n} \cdot \hat{e}_z)$. $X_r$ is planar. A circle or ellipse is fitted to the $x_i$ and $y_i$ components of $X_r$ (see for instance Suppl. section 8.2, 8.4).
Finally, the centres $\bar{x} = (x_c, y_c, z_c)$ of the fitted circle and ellipse are shifted and Rodrigues-rotated back to match the non-planar $X$ according to

$$\vec{x}_c = \bar{x} + R(\hat{k}, \theta)P. \tag{42}$$

The rotation matrix $R$ is derived from (41). Here, $P = (x_c, y_c, 0)$, the axis of rotation is $\hat{k} = (\hat{e}_z \times \hat{n}^T)/||\hat{e}_z \times \hat{n}^T||$ and the angle of rotation is $\theta = \arccos(\hat{e}_z \cdot \hat{n})$.
A 3D-fitted circle is fully characterised by radius $r$, $\vec{x}_c$, $\hat{t}_1$, and $\hat{t}_2$.
A vertex $\vec{x}_v = (x_v, y_v)$ and co-vertex $\vec{x}_{cv} = (x_{cv}, y_{cv})$ of the fitted ellipse (Suppl. section 8.2, (55)) are the end-points of a semi-major and semi-minor axis respectively and are not necessarily collinear to $\hat{t}_1$ and $\hat{t}_2$. The vectors $\vec{x}_v$ and $\vec{x}_{cv}$ are therefore Rodrigues-rotated to match $X$ following

$$\vec{x}'_{v,vc} = R(\hat{k}, \theta)P. \tag{43}$$

With $P = (x_v, y_v, 0)$ or $P = (x_{cv}, y_{cv}, 0)$, $\hat{k} = (\hat{e}_z \times \hat{n}^T)/||\hat{e}_z \times \hat{n}^T||$ and $\theta = \arccos(\hat{e}_z \cdot \hat{n})$.
An 3D-fitted ellipse is fully characterised by $\vec{x}_c$, $\vec{x}'_v$, and $\vec{x}'_{cv}$.

#### 8.2 Fitting an Ellipse

A standard ellipse has a centre $(0, 0)$, and axes $b$ and $a$ that align with the x and y axes of a Cartesian coordinate system following

$$\frac{x^2}{a^2} + \frac{y^2}{b^2} = 1. \tag{44}$$

This ellipse can be rotated anticlockwise by $\rho$ and translated to a new centre $(x_c, y_c)$, expanding (44) into

$$\frac{((x - x_c)\cos(\rho) + (y - y_c)\sin(\rho))^2}{a^2} + \frac{((x - x_c)\sin(\rho) - (y - y_c)\cos(\rho))^2}{b^2} = 1. \tag{45}$$

This can be written in polynomial form and the coefficients of the equation can substituted with $A$ to $F$ to yield

$$
\begin{aligned}
&AX^2 + BXY + CY^2 + DX + EY + F = 0,\\
&\text{with}\\
&A = a^2 \sin^2(\rho) + b^2 \cos^2(\rho),\\
&B = 2(b^2 - a^2) \sin(\rho) \cos(\rho),\\
&C = a^2 \cos^2(\rho) + b^2 \sin^2(\rho),\\
&D = -2Ax_c - By_c,\\
&E = -Bx_c - 2Cy_c,\\
&F = Ax_c^2 + Bx_c y_c + Cy_c^2 - a^2 b^2.
\end{aligned} \tag{46}
$$

Assuming $F = -1$, this yields

$$
AX^2 + BXY + CY^2 + DX + EY = 1. \tag{47}
$$

A matrix $M$ is constructed from the $x, y$ coordinates of model NPC nodes, whereby the columns represent the indeterminates of (47) and the rows represent the nodes $1 \ldots n$. Depending on which part of the NPC the ellipse is fitted to, $n$ is either all nodes of model NPC ($n \in 1 \ldots m$), or all nodes within the CR or NR, or all nodes within a ring ($n \in 1 \ldots s$).

$$
M = \begin{bmatrix} x_1^2 & x_1 y_1 & y_1^2 & x_1 & y_1 \\ \vdots & \vdots & \vdots & \vdots & \vdots \\ x_n^2 & x_n y_n & y_n^2 & x_n & y_n \end{bmatrix} \tag{48}
$$

The coefficients are written in column vector form $\vec{b}$ as

$$
\vec{b} = \begin{bmatrix} A \\ B \\ C \\ D \\ E \end{bmatrix}, \tag{49}
$$

and optimised using least squares with

$$
\arg\min_b \left\| M\vec{b} - \vec{1} \right\|^2. \tag{50}
$$

The values of $a$ and $b$ are then determined with

$$
a, b = \frac{-\sqrt{2(AE^2 + CD^2 - BDE + (B^2 - 4AC)F)((A + C) \pm \sqrt{(A - C)^2 + B^2})}}{B^2 - 4AC}. \tag{51}
$$

In (51), either of $a$ or $b$ can corresponds to the semimajor axis. We therefore redefine $a'$ as semimajor, and $b'$ as semiminor:

$$
\begin{aligned}
a' &= \max(a, b),\\
b' &= \min(a, b).
\end{aligned} \tag{52}
$$

Furthermore, we determine the rotation $\rho$ as

$$
\rho = \begin{cases} \arctan(\frac{1}{B}(C - A - \sqrt{(A - C)^2 + B^2}) & \text{for } B \neq 0, \\ 0 & \text{for } B = 0, A < C, \\ 0.5\pi & \text{for } B = 0, A > C. \end{cases} \tag{53}
$$

Next, $\rho$ is expressed as a value between $[-0.5\pi, 0.5\pi]$ with

$$\rho' = \begin{cases} \rho - 0.5\pi & \text{if } A < 0 \text{ and if } \rho \geq 0, \\ \rho + 0.5\pi & \text{if } A < 0 \text{ and if } \rho < 0. \end{cases}$$
$$\text{and} \tag{54}$$
$$\rho = \begin{cases} \rho' & \text{if } A < 0, \\ \rho & \text{otherwise.} \end{cases}$$

Consequently, $\rho$ is the angle the right-hand semimajor axis forms with respect to a horizontal line.
The vertices and co-vertices of the ellipse are the points at the end of the major and minor axes respectively. A vertex $\vec{x}_v$ and co-vertex $\vec{x}_{cv}$ can be determined via

$$\begin{aligned} \vec{x}_v &= a'(\cos(\rho), \sin(\rho)), \\ \vec{x}_{cv} &= b'(\cos(\rho - 0.5\pi), \sin(\rho - 0.5\pi)), \end{aligned} \tag{55}$$

#### 8.3 Finding the RSS

We fit a circle based on least-squares [11], the RSS is thereby minimised, thus can be directly read out. As circle and ellipse are fit to planar projections $X_r$ of the NPC (41), this RSS only reflects the lateral direction and is henceforth called $\text{RSS}_l$.
As $X_r$ is planar and zero-centred, the residual sum of squares in axial direction ($\text{RSS}_{ax}$) from the fitted circle or ellipse can be obtained by summing the squared z-coordinates $z_{r,i}$ in $X_r$ For both circle-and ellipse fit, the overall RSS is the sum of its axial- and lateral components.

$$\text{RSS}_{ax} = \sum_{i=1}^{m} z_{r,i}^2 \tag{56}$$

$$\text{RSS} = \text{RSS}_{ax} + \text{RSS}_l \tag{57}$$

#### 8.4 Approximating the lateral RSS of the fitted ellipse

As for the fitted circle, the RSS of nodes to the fitted ellipse is a measure of deviation from it. The $\text{RSS}_{ax}$ in axial direction of the ellipse-plane can be obtained via (56). However, the fitting procedure described in Suppl. section 8.2 does not straight-forwardly return the lateral sum of squared Euclidean distances $\text{RSS}_l$ of nodes to the ellipse. We therefore estimate this measure by approximating the closest point on the fitted ellipse for each node [12], and then summing up the squared Euclidean distances of all pairs of nodes and closest points.
The model NPC and fitted ellipse are first transformed into standard ellipse form. For this, the model NPC is rotated by $-\rho$ around the centre of the fitted ellipse $(x_c, y_c)$ and then translated so that its centre of mass is at $(0, 0)$. It consequently aligns with a standard ellipse with semimajor axis $a'$ and semiminor axis $b'$ of the fitted ellipse, henceforth called $a$ and $b$. Therefore, the $\text{RSS}_l$ between this rotated and shifted model NPC to the standard ellipse with $a, b$ is equivalent to the $\text{RSS}_l$ between the model NPC and the fitted ellipse. Furthermore, projecting all $m$ nodes with coordinates $\vec{x} = (x_i, y_i)$, $i \in 1 \dots m$ onto the 1st quadrant ($x'_i = |x_i|, y'_i = |y_i|$, $\vec{x}' = (x'_i, y'_i)$) does not change the $\text{RSS}_l$. Any point on the ellipse can be approximated with a circle of the same curvature, whereby the centre of all such circles form a curve called evolute. The closest point on the ellipse $\vec{p}_i = (p_{x_i}, p_{y_i})$ of any point $\vec{x}' = (x'_i, y'_i)$ is therefore the tangential point between a circle around $\vec{x}'$ and the

ellipse, or alternatively the circle approximating the ellipse at this point with corresponding centre $\vec{e}_i = (e_{xi}, e_{yi})$ on the evolute.
Both the ellipse and evolute can be parameterized based on an angle $\tau$ following

$$\begin{aligned} p_x &= a\cos(\tau), \\ p_y &= b\sin(\tau), \end{aligned} \tag{58}$$

$$\begin{aligned} e_x &= (a^2 - b^2)\frac{\cos(\tau)^3}{a}, \\ e_y &= (b^2 - a^2)\frac{\sin(\tau)^3}{b}. \end{aligned} \tag{59}$$

Consequently, for a point $\vec{x}'_i$, $\tau$ for the closest point $\vec{p}$ needs to be approximated, which is done iteratively:

- As all nodes have been projected to the 1st quadrant, $\tau$ is initialised as the bisector of the 1st quadrant, $\tau_0 = \pi/4$ rad.
- The corresponding point on the ellipse is $\vec{p}(\tau_0) = (p_x(\tau_0), p_y(\tau_0))$, whereby $p_x(\tau_0) = a\cos(\tau_0)$ and $p_y(\tau_0) = b\sin(\tau_0)$.
- The centre of curvature of $\vec{p}(\tau_0)$ is at $\vec{e}(\tau_0)$, which is computed analogously using (59).
- The radius of curvature at $\tau_0$ is therefore $||\vec{r}(\tau_0)||$ with $\vec{r}(\tau_0) = (r_x(\tau_0), r_y(\tau_0)) = (p_x(\tau_0) - e_x(\tau_0), p_y(\tau_0) - e_y(\tau_0))$.
- Furthermore, the vector $\vec{q}(\tau_0)$ connects $\vec{e}(\tau_0)$ with $\vec{x}'_i$: $\vec{q}(\tau_0) = (q_x(\tau_0), q_y(\tau_0)) = (x'_i - e_x(\tau_0), y'_i - e_y(\tau_0))$.
- The arc length $\triangle c(\tau_0)$ between $\vec{r}(\tau_0)$ and $\vec{q}(\tau_0)$ on a circle around $\vec{e}(\tau_0)$ with radius $||\vec{r}(\tau_0)||$ is given by

$$\begin{aligned} &\triangle c(\tau_0) = ||\vec{r}||(\tau_0)\arcsin\left(\frac{||\vec{r}(\tau_0) \times \vec{q}(\tau_0)||}{||\vec{r}||(\tau_0)||\vec{q}||(\tau_0)}\right), \\ &\text{with} \\ &\vec{r}(\tau_0) \times \vec{q}(\tau_0) = r_x(\tau_0)q_y(\tau_0) - r_y q_x(\tau_0). \end{aligned} \tag{60}$$

- Following, a new $\tau_1$ is found that approximately minimises $\triangle c(\tau_0)$ with

$$\triangle\tau = \frac{\triangle c(\tau_0)}{\sqrt{a^2 + b^2 - (p_x(\tau_0))^2 - (p_y(\tau_0))^2}}, \tag{61}$$

$$\tau_1 = \tau_0 + \triangle\tau. \tag{62}$$

- Next, $\tau_1$ is ensured to remain in the 1st quadrant with

$$\begin{aligned} &\tau'_1 = \min(0.5\pi, \max(0, \tau_1)), \\ &\tau_1 = \tau'_1. \end{aligned} \tag{63}$$

- The variable $\tau_1$ updates $\vec{e}(\tau_1)$ and $\vec{p}(\tau_1)$.

The value of $\triangle_c(\tau_0)$ is the arc-length on the circle that approximates curvature at $\vec{p}(\tau_0)$, not on the ellipse. Therefore $\vec{p}(\tau_1)$ might not correspond to the closest point to $\vec{x_i'}$. However, when the above stated procedure is repeated for $j \in 0 \ldots n$ iterations, the circle approximating curvature at $\vec{p}(\tau_j)$ increasingly resembles the ellipse at the closest point to $\vec{x}'$. We set $n$ to 2 and define $\vec{p}(\tau_2)$ as the closest point to $\vec{x_i'}$. We then project $\vec{p}(\tau_2)$ onto the original quadrant of node $\vec{x}_i$ as $p_x = p_x(\tau_2)\text{sgn}(x_i), p_y = p_y(\tau_2)\text{sgn}(y_i)$. Henceforth, we call the closest point $\vec{p}$ to $\vec{x}_i$ "$\vec{p}_i$".
Finally, the $\text{RSS}_l$ is computed with

$$\text{RSS}_l = \sum_{i=1}^{n} (\vec{x}_i - \vec{p}_i)^2, \tag{64}$$

where $n$ is the number of nodes.
For a circular NPC, this $\text{RSS}_l$ is almost the same as for the fitted circle. For an elongated NPC, this $\text{RSS}_l$ is smaller than that of a fitted circle. When all nodes are close to the ellipse, the $\text{RSS}_l$ is close to 0. The 3D RSS is finally obtained via (57).

### 9 Additional features

#### 9.1 Ring distance and ring shift

Ellipses are fitted to both CR and NR and have centres $\vec{x}_{c,\text{CR}} = (c_{\text{CR},x}, c_{\text{CR},y}, c_{\text{CR},z})$ and $\vec{x}_{c,\text{NR}} = (c_{\text{NR},x}, c_{\text{NR},y}, c_{\text{NR},z})$ respectively. The ring distance of deformed NPCs is here defined as $d = c_{\text{CR},z} - c_{\text{NR},z}$. The shift is defined as shift $= ||(c_{\text{CR},x}, c_{\text{CR},y}) - (c_{\text{NR},x}, c_{\text{NR},y})||$.

#### 9.2 Tilt angles

The normal unit vector indicating tilt of the CR-plane $\hat{n}_{\text{CR}}$ is determined by fitting a 3D ellipse. The corresponding normal unit vector for the NR-plane is given by $\hat{n}_{\text{NR}}$. Vectors $\hat{n}_{\text{CR}}$ and $\hat{n}_{\text{NR}}$ form an angle $\alpha_{\text{tilt}} = \arccos(\hat{n}_{\text{CR}} \cdot \hat{n}_{\text{NR}})$, indicating the tilt of CR and NR relative to each other. If $\alpha_{\text{tilt}} > 0.5\pi$, it is updated to $\alpha'_{\text{tilt}} = \pi - \alpha_{\text{tilt}}$.

### 10 Output files

Simulated NPCs are saved as CSV files that can be loaded into photophysics simulation software (e.g. [13]). Each node is described by coordinates x, y, z and an index per NPC. We do not introduce further rotation, translation or tilt to the NPC, as most downstream photophysics simulation software includes this functionality.

### 11 Included NPC models

All thus far selectable NPC models are indicated in Table S2.

### 12 Simulated Nups

The N-terminus (N-term.) and C-terminus (C-term.) respectively denote the first and last structurally resolved amino acid, whose coordinates are used for simulations. Value n in CR, NR, IR, and in bridge denote the number of rings in each. The bridge is a linker between the OR and IR Table S3.

**Table S2.** Input models

| Model | Substructure | Origin | Resolution[1](Å) |
|---|---|---|---|
| 5A9Q [3] | ORs | HeLa, isolated NE | 23 |
| 7PEQ [1] | ORs | DLD-1, native environment | 35 |
| 5IJN [14] | IR | HeLa, isolated NE | 21.4 |
| 5IJO [14] | IR | HeLa, isolated NE | 21.4 |
| 7PER [1] | IR | DLD-1, native environment | 35 |
| 7R5K [2] | Whole scaffold | HeLa, isolated NE | 12 |
| 7R5J [2] | Whole scaffold | HeLa, HEK293, native environment | 50 |

[1] As reported in PDB.

**Table S3.** Input models in detail

| Model | Nup | N-term. | C-term. | n in CR | NR | IR | bridge | total |
|---|---|---|---|---|---|---|---|---|
| 5A9Q | Nup43 | 4 | 380 | 2 | 2 | | | 4 |
| | Nup160 | 41 | 1201 | 2 | 2 | | | 4 |
| | Nup37 | 9 | 326 | 2 | 2 | | | 4 |
| | Sec13 | 14 | 304 | 2 | 2 | | | 4 |
| | Seh1 | 1 | 322 | 2 | 2 | | | 4 |
| | Nup85 | 9 | 475 | 2 | 2 | | | 4 |
| | Nup155 | 20 | 730 | | | | 2 | 2 |
| | Nup133 | 518 | 1156 | 2 | 2 | | | 4 |
| | Nup96 | 277 | 751 | 2 | 2 | | | 4 |
| | Nup107 | 150 | 924 | 2 | 2 | | | 4 |
| 7PEQ | Nup133 | 518 | 1156 | 2 | 2 | | | 4 |
| | Nup107 | 150 | 924 | 2 | 2 | | | 4 |
| | Nup96 | 333 | 922 | 2 | 2 | | | 4 |
| | Sec13 | 14 | 304 | 2 | 2 | | | 4 |
| | Seh1 | 5 | 320 | 2 | 2 | | | 4 |
| | Nup85 | 20 | 651 | 2 | 2 | | | 4 |
| | Nup43 | 4 | 380 | 2 | 2 | | | 4 |
| | Nup160 | 78 | 1195 | 2 | 2 | | | 4 |
| | Nup37 | 18 | 324 | 2 | 2 | | | 4 |
| 7PER | Nup205 | 9 | 1692 | | | 4 | | 4 |
| | Nup155 | 20 | 1375 | | | 4 | | 4 |
| | Nup93 | 173 | 815 | | | 4 | | 4 |
| | Nup54 | 128 | 493 | | | 4 | | 4 |
| | Nup58[1] | 248 | 418 | | | 4 | | 4 |
| | Nup62 | 334 | 502 | | | 4 | | 4 |
| 5IJN | Nup205 | 9 | 1692 | | | 4 | | 4 |
| | Nup54 | 128 | 493 | | | 4 | | 4 |
| | Nup58 | 248 | 418 | | | 4 | | 4 |
| | Nup155 | 20 | 2 x 863<br>4 x 1375 | | | 4 | 2 | 6 |
| | Nup93 | 1 | 815 | | | 4 | | 4 |
| | Nup62 | 334 | 502 | | | 4 | | 4 |
| 5IJO | Nup54 | 128 | 493 | | | 4 | | 4 |
| | Nup188 | 1 | 1564 | | | 2 | | 2 |
| | Nup205 | 9 | 1692 | | | 2 | | 2 |
| | Nup155 | 20 | 2 x 863<br>4 x 1375 | | | 4 | 2 | 6 |
| | Nup93 | 1 | 815 | | | 4 | | 4 |

| | | | | | | | | |
|---|---|---|---|---|---|---|---|---|
| | Nup58[1] | 248 | 418 | | | 4 | | 4 |
| | Nup62 | 334 | 502 | | | 4 | | 4 |
| 7R5J 7R5K | RANBP2 | 4 | 759 | 5 | | | | 5 |
| | Nup210[2] | 2 | 1832 | | | 8 | | 8 |
| | ALADIN | 2 | 486 | | | 2 | | 2 |
| | Nup93 | 2 | 819 | | | 4 | | 4 |
| | Nup93 | 94 | 819 | 2 | 1 | | | 3 |
| | Nup188 | 2 | 1749 | | | 2 | | 2 |
| | Nup205 | 2 | 2012 | 2 | 1 | 2 | | 5 |
| | Nup155 | 10 | 1391 | | | 4 | 2 | 6 |
| | NDL1 | 2 | 674 | | | 2 | | 2 |
| | Nup35 | 86 | 326 | | | 4 | | 4 |
| | Nup54 | 111 | 493 | | | 4 | | 4 |
| | Nup58[1] | 246 | 418 | | | 4 | | 4 |
| | Nup62 | 332 | 502 | 1 | | 4 | | 5 |
| | Nup133 | 71 | 1156 | 2 | 2 | | | 4 |
| | Nup107 | 144 | 925 | 2 | 2 | | | 4 |
| | Nup96 | 232 | 937 | 2 | 2 | | | 4 |
| | SEC13 | 2 | 302 | 2 | 2 | | | 4 |
| | SEH1 | 2 | 324 | 2 | 2 | | | 4 |
| | Nup85 | 2 | 656 | 2 | 2 | | | 4 |
| | Nup43 | 2 | 380 | 2 | 2 | | | 4 |
| | Nup160 | 38 | 1436 | 2 | 2 | | | 4 |
| | Nup37 | 5 | 326 | 2 | 2 | | | 4 |
| | ELYS | 2 | 1005 | | 2 | | | 2 |
| | Nup98 | 731 | 880 | | 1 | | | 1 |
| | Nup98 | 597 | 615 | 2 | | 4 | | 6 |
| | Nup214 | 700 | 972 | 1 | | | | 1 |
| | Nup88 | 7 | 741 | 1 | | | | 1 |

[1] Reported as p58/p45
[2] Primarily in NE lumen

**Table S4.** ChimeraX commands to extract RU coordinates

| Model | Nup | Terminus | Command |
|---|---|---|---|
| 5A9Q | Nup43 | N | getcr #1.1/0,R,9,I:4@CA |
| | | C | getcr #1.1/0,R,9,I:380@CA |
| | Nup160 | N | getcr #1.1/1,S,a,J:41@CA |
| | | C | getcr #1.1/1,S,a,J:1201@CA |
| | Nup37 | N | getcr #1.1/2,T,b,K:9@CA |
| | | C | getcr #1.1/2,T,b,K:326@CA |
| | Sec13 | N | getcr #1.1/6,X,F,O:14@CA |
| | | C | getcr #1.1/6,X,F,O:304@CA |
| | Seh1 | N | getcr #1.1/7,Y,G,P:1@CA |
| | | C | getcr #1.1/7,Y,G,P:322@CA |
| | Nup85 | N | getcr #1.1/Q,8,Z,H:9@CA |
| | | C | getcr #1.1/Q,8,Z,H:475@CA |
| | Nup155 | N | getcr #1.1/A,B:20@CA |
| | | C | getcr #1.1/A,B:730@CA |
| | Nup133 | N | getcr #1.1/3,U,C,L:518@CA |
| | | C | getcr #1.1/3,U,C,L:1156@CA |
| | Nup96 | N | getcr #1.1/5,W,E,N:277@CA |

| | | | |
|---|---|---|---|
| | | C | getcr #1.1/5,W,E,N:751@CA |
| | Nup107 | N | getcr #1.1/M,D,V,4:150@CA |
| | | C | getcr #1.1/M,D,V,4:924@CA |
| | Nup133 | N | getcrd /?C:518@CA |
| | | C | getcrd /?C:1156@CA |
| | Nup107 | N | getcrd /?D:150@CA |
| | | C | getcrd /?D:924@CA |
| | Nup96 | N | getcrd /?E:333@CA |
| | | C | getcrd /?E:922@CA |
| | Sec13 | N | getcrd /?F:14@CA |
| | | C | getcrd /?F:304@CA |
| 7PEQ[1] | Seh1 | N | getcrd /?G:5@CA |
| | | C | getcrd /?G:320@CA |
| | Nup85 | N | getcrd /?H:20@CA |
| | | C | getcrd /?H:651@CA |
| | Nup43 | N | getcrd /?I:4@CA |
| | | C | getcrd /?I:380@CA |
| | Nup160 | N | getcrd /?J:78@CA |
| | | C | getcrd /?J:1195@CA |
| | Nup37 | N | getcrd /?K:18@CA |
| | | C | getcrd /?K:324@CA |
| | Nup205 | N | getcrd /D,J,V,P:9@CA |
| | | C | getcrd /D,J,V,P:1692@CA |
| | Nup155 | N | getcrd /E,K,Q,W:20@CA |
| | | C | getcrd /E,K,Q,W:1375@CA |
| | Nup93 | N | getcrd /C,I,O,U:173@CA |
| | | C | getcrd /C,I,O,U:815@CA |
| 7PER[1] | Nup54 | N | getcrd /F,L,R,X:128@CA |
| | | C | getcrd /F,L,R,X:493@CA |
| | Nup58 "P58P45" | N | getcrd /G,M,S,Y:248@CA |
| | | C | getcrd /G,M,S,Y:418@CA |
| | Nup62 | N | getcrd /H,N,T,Z:334@CA |
| | | C | getcrd /H,N,T,Z:502@CA |
| | Nup205 | N | getcrd #1.1/D,J,P,V:9@CA |
| | | C | getcrd #1.1/D,J,P,V:1692@CA |
| | Nup54 | N | getcrd #1.1/F,L,R,X:128@CA |
| | | C | getcrd #1.1/F,L,R,X:493@CA |
| | Nup58 | N | getcrd #1.1/G,M,S,Y:248@CA |
| | | C | getcrd #1.1/G,M,S,Y:418@CA |
| 5IJN | Nup155 | N | getcrd #1.1/A,B,E,K,Q,W:20@CA |
| | | C | getcrd #1.1/A,B:863@CA |
| | | | getcrd #1.1/E,K,Q,W:1375@CA |
| | Nup93 | N | getcrd #1.1/C,I,O,U:1@CA |
| | | C | getcrd #1.1/C,I,O,U:815@CA |
| | Nup62 | N | getcrd #1.1/H,N,T,Z:334@CA |
| | | C | getcrd #1.1/H,N,T,Z:502@CA |
| 5IJO | Nup54 | N | getcrd /F,L,R,X:128@CA |
| | | C | getcrd /F,L,R,X:493@CA |
| | Nup188 | N | getcrd /J,V:1@CA |
| | | C | getcrd /J,V:1564@CA |
| | Nup205 | N | getcrd /D,P:9@CA |
| | | C | getcrd /D,P:1692@CA |

| | | |
|---|---|---|
| Nup155 | N | getcrd /A,B,E,K,Q,W:20@CA |
| | C | getcrd /A,B:863@CA<br>getcrd /E,K,Q,W:1375@CA |
| Nup93 | N | getcrd /C,I,O,U:1@CA |
| | C | getcrd /C,I,O,U:815@CA |
| Nup58 "P58P45" | N | getcrd /G,M,S,Y:248@CA |
| | C | getcrd /G,M,S,Y:418@CA |
| Nup62 | N | getcrd /H,N,T,Z:334@CA |
| | C | getcrd /H,N,T,Z:502@CA |

1 Command for PDB file with single RU. Output verified on Biological Assembly with all RUs (not shown).

### 13 Photophysics simulation parameters

**Table S5.** Photophysics simulation parameters for Nup107 dataset, modified from Wu et al., 2022 [15]

| SMLM property | value | units/notes |
|---|---|---|
| Photon count | 12000 | photons/pixel/localization |
| Background | 140 | |
| Labelling efficiency | 67 | % |
| Re-activation | 4.1 | |
| Lifetime | 1 | frames |
| Linkage error (free) | 3.2 | nm from [16] |
| EM on | True | simulates EMCCD sensor |
| Number of frames | 100000 | |

**Table S6.** Photophysics simulation parameters for Nup96 dataset

| SMLM property | value | units/notes |
|---|---|---|
| Photon count[1] | 10150 ± 1000 | photons/pixel/localization |
| Background | 260 | From [17]. More noisy than 140 [15] |
| Labelling efficiency[1] | 58 | % |
| Re-activation[1] | 2.9 | |
| Lifetime[2] | 2 | frames |
| Linkage error (free) | 3.2 | nm, from [16] |
| EM on[3] | True | simulates EMCCD sensor |
| Number of frames[4] | 60000 | |
| Random rot theta | True, 15 | Degrees. Tilts whole NPC as nucleus not always flat |
| Rendering: Locprec[5] | 0-10 | nm |

1 Thevathasan et al. [18], Table 1, 35 mM MEA + GLOX buffer
2 By comparing Thevathasan et al. [18] Supplementary Table 4: Frametime 30/40 ms, intensity 6 $kW/cm^2$ and Wu et al. [15] Methods, Microscope setup and imaging: Framerate 100 ms, intensity 6 $kW/cm^2$, Simulated lifetime 1.
3 Thevathasan et al. [18], Methods, Microscopy. Microscope setup and imaging: Evolve512D EMCCD camera (Photometrics).
4 From [18], Supplementary Table 4.
5 Thevathasan et al. [18] Supplementary: Fitting of raw data in SMAP, Rendering of localizations
